# Metabolic cost as a determinant of light quality acclimation: a full-PAR characterization of the CA3 cyanobacterium *Nostoc* sp. CCAP 1453/38

**DOI:** 10.64898/2026.07.27.740936

**Authors:** Tomáš Zavřel, Anne-Christin Pohland, Mariann Kis, Martin Lukeš, Anna Segečová, László Kovács, Jan Mareš, Zoltán Novák, Jan Červený, Gábor Bernát

## Abstract

Light quality acclimation is one of the key drivers of cyanobacterial physiology, ecology, and productivity. Although chromatic acclimation type 3 (CA3) is the canonical example of light quality–driven phycobilisome remodeling, full-PAR physiological characterization of CA3 strains has so far been limited. Here, we characterize the spectral acclimation strategies of the cyanobacterium *Nostoc* sp. CCAP 1453/38 across the full photosynthetically active radiation (PAR) range. Genomic analysis confirmed *Nostoc* as a CA3 strain capable of dynamically adjusting phycoerythrin (PE) and phycocyanin (PC) content in its phycobilisome (PBS) rods. During cultivation under narrow-band LEDs, PE was upregulated under violet, blue and green light (435–555 nm), optimizing light harvesting primarily in the blue-green part of the PAR spectrum, while PC was upregulated under red light (633–687 nm). Beyond canonical CA3 pigment switching, *Nostoc* responded to wavelengths poorly absorbed by PBS in two qualitatively different ways. Under growth-constraining blue light (465 nm), the strain upregulated total PBS and photosystem II (PSII) levels and biased phycobilisome coupling toward PSII (an increased PBS–PSII/PBS–PSI ratio). However, this metabolically costly response could not overcome the underlying excitonic imbalance caused by preferential PSI excitation, resulting in a low cell division rate. Under near far-red light (687 nm), PBS–PSII coupling itself was enhanced, yet total PBS content was reduced rather than increased. Specific growth rates remained as high as under red light, suggesting that this PBS–PSII reorganization avoided the metabolic burden of antenna upregulation. These results indicate that the same underlying challenge of PSII under-excitation can trigger qualitatively different acclimation responses, only some of which are energetically affordable. Spectral acclimation thus depends not only on matching pigment composition to incident wavelengths, but also on the metabolic cost of the response. Compared with parallel datasets on CA1 and non-CA strains obtained under identical conditions, our findings extend CA3 characterization beyond the canonical green/red framework, highlight bottlenecks and advantages of light quality acclimation in *Nostoc*, and provide a physiological basis for optimizing light regimes in controlled cyanobacterial cultivations.

## 1. Introduction

About 50% of the total primary production on Earth is attributable to phytoplankton (Carvalho et al., 2017; Mattei and Scardi, 2021), and cyanobacteria alone account for about 10-25% (Flombaum et al., 2013). These substantial contributions strongly depend on the ability of these cells to acclimate to the prevailing environmental conditions. In aquatic environments, the phytoplankton fitness and distribution are determined by a variety of factors, including temperature (Righetti et al., 2019), nutrient ratios and concentrations (Martiny et al., 2013), pH (Hyun et al., 2020), salinity (Sugie et al., 2020), water movement and mixing (Wirtz and Smith, 2021), turbidity (Cloern, 1987), presence or absence of grazers (Matsumoto et al., 2026) and/or toxic substances (Zitoun et al., 2024), and, notably, light - the primary energy source for all phototrophs. During photosynthesis, solar energy is captured by light-harvesting pigment-protein complexes – such as phycobilisome (PBS) in cyanobacteria – and the excitonic energy is funneled to the reaction centers of photosystem II (PSII) and photosystem I (PSI). The excitons undergo charge separation in the PSII and PSI reaction centers, driving both linear and cyclic electron flows along the photosynthetic electron transport chain (PETC). Ultimately, these processes generate energy (i.e. ATP) and reducing equivalents (NADPH) that fuel the fixation of inorganic carbon into organic molecules through the Calvin-Benson-Bassham (CBB) cycle.

Light availability changes with the depth of the water column (Stomp et al., 2007). It has been recognized that it is not only the absolute amount of photons available to phytoplankton cells, but also the light quality that shapes phytoplankton biogeography (Mattei et al., 2025). The underwater light spectrum differs substantially from the sunlight spectrum available to plants in the atmosphere. Due to the specific vibrational modes of water molecules, and to the presence of humic and/or inorganic matter in the water column, the underwater spectrum can be dominated either by blue, green, orange, red or even far red photons (Holtrop et al., 2021). These specific color niches can provide fitness advantages to certain phytoplankton strains, such as the so-called blue-light- or green-light-specialist strains of marine *Synechococcus* (Dufour et al., 2025). The ability to harvest light in a narrow spectral range strongly depends on the type of pigmentation. Blue-light-specialists within the *Synechococcus* genus possess phycoerythrin complexes with a high ratio of the blue light-absorbing chromophore phycourobilin (PUB) to the green light-absorbing chromophore phycoerythrobilin (PEB). On the contrary, green light specialists possess low PUB:PEB ratios (Palenik, 2001). Another chromophore present in PBS, phycocyanobilin (PCB), allows efficient harvesting of red light. (Scheer and Zhao, 2008).

PBSs are typically composed of a central core and radiating rods, both of which form cylindrical structures constituted by phycobiliproteins and associated linker proteins. Phycobiliproteins are classified into four types: phycocyanin (PC), phycoerythrin (PE), phycoerythrocyanin (PEC), and allophycocyanin (APC) (Watanabe and Ikeuchi, 2013). These classes differ in their covalently bound bilins, which set their absorption maxima: APC and PC carry phycocyanobilin (PCB), absorbing red light (∼650 nm and ∼620 nm, respectively); PE carries PEB (∼560 nm) and, in some strains PUB (∼495 nm), shifting absorption toward green and blue-green; and PEC carries PVB (∼570 nm) on its α-subunit together with PCB on its β-subunit (Scheer and Zhao, 2008). The phycobiliproteins assemble into disc-shaped units that, when connected by linker proteins, enable the unidirectional transfer of captured light energy through the phycobilisome – from PE to PC (or PEC, where present) within the rods, and subsequently to APC in the core – ultimately delivering energy to PSII (Chang et al., 2015) or PSI (van Stokkum et al., 2023).

Cyanobacteria have also evolved molecular mechanisms to efficiently harvest light across multiple spectral niches. The structure and pigment composition of the PBS can be adjusted according to the quality of the ambient light through the process of chromatic acclimation (CA). To date, eight different types of CA (CA0-CA7) have been identified; a single cyanobacterium can possess either one or, in some cases, two of these types (Hirose et al., 2019; Sanfilippo et al., 2019). Specifically, CA0 and CA7 optimize light absorption in the yellow-green spectral range by regulating the structure and docking of rods *via* the CpcL linker protein (CA0) or by adjusting the PEC content (CA7) (Hirose et al., 2019). During CA1, red light promotes the formation of canonical PBS containing the rod-core linker CpcG1, which anchors the rods to the APC core. In contrast, green light induces, again, the synthesis of the alternative linker CpcL, resulting in the assembly of PBS that lack core components and preferentially associate with PSI. During CA2, absorption of green light is optimized by the upregulation of PE, leading to the extension of PBS rods, while the amount of PC remains unchanged. CA3 coordinates the complementary upregulation of PE and PC under green and red light, respectively, by adjusting both protein and chromophore content for broader spectral coverage. In CA4, adaptation to blue or green light is achieved through changes in the PUB to PEB ratio, without modifying the protein composition of the PBS. CA5 and CA6 are associated with far-red light acclimation. In CA5, PC-containing PBSs are replaced by chlorophyll *d*-based light harvesting complexes in the thylakoid membrane. In CA6, far-red light induces a more complex response, including the expression of far-red–absorbing APC, PSII and PSI, as well as a shift from chlorophyll *a* (Chl *a*) to chlorophylls *d* or *f* – a process known as FaRLiP (far-red light photoacclimation). Despite CA3 being the first described chromatic acclimation type historically and the basis of the term “complementary chromatic adaptation”, the physiological characterization of CA3 strains has thus far been confined almost entirely to *Fremyella diplosiphon*, and to comparisons under only two wavelengths: green and red. The behavior of CA3 strains across the full PAR spectrum, and how it compares quantitatively to non-CA strains under the same conditions, remains essentially unexplored.

Besides CA, light harvesting can be optimized by tuning the PSII/PSI stoichiometry (Bernát et al., 2021), state transitions (Calzadilla and Kirilovsky, 2020), thermal dissipation of excess excitation energy via orange carotenoid protein-mediated non-photochemical quenching (OCP-NPQ) (Kirilovsky and Kerfeld, 2016), excitonic decoupling of the PBS (Tamary et al., 2012), activation of futile cycles (Tchernov et al., 2003; Müller et al., 2019), or shifts in transcriptome (Luimstra et al., 2020), proteome (Zavřel et al., 2019), or lipids (Zavřel et al., 2018b). Many of these acclimations correlate with shifts in the redox state of the plastoquinone (PQ) pool (Bernát et al., 2021; Zavřel et al., 2024).

This work describes the light acclimation of *Nostoc* sp. CCAP 1453/38 (referred to as *Nostoc* hereafter), a CA3-type cyanobacterium, across the full PAR spectrum (400-700 nm). The experimental setup, used in several previous studies, allows for a qualitative comparison of the photoacclimation traits with the CA1 cyanobacterium *Synechocystis* sp. PCC 6803 (Zavřel et al., 2024) and with CA-incapable strains such as *Cyanobium gracile* and *Cyanobium* sp. NIVA-CYA 375 (Bernát et al., 2021; Kis et al., 2026). The comparative analysis reveals a growth advantage of PE-rich strains under green light (495-555 nm). However, additional light quality acclimation characteristics, unique to each strain, were identified across the PAR spectrum. These include shifts in the PSII/PSI ratio, PBS attachment to PSII or PSI, linear and cyclic electron flows within PETC, the redox state of the PQ pool, and total PBS content. These differences show distinct acclimation strategies to optimize light-harvesting not only under unfavorable light (blue or near far-red light) but also under optimal light conditions.

In this study, we demonstrate that *Nostoc* sp. CCAP 1453/38 utilizes wavelength-specific strategies beyond simple pigment switching. While CA3-driven PE upregulation optimizes growth under green light, a significant bioenergetic bottleneck occurs under blue light due to the metabolic cost of PBS antenna upregulation. Conversely, near far-red light triggers efficient photosystem reorganization without additional pigment synthesis. These findings reveal the fundamental energetic trade-offs governing spectral plasticity and ecological success in CA3-type cyanobacteria.

## 2. Materials and Methods

### 2.1 Inoculum cultures and experimental setup

The strain *Nostoc* sp. CCAP 1453/38 was originally isolated from a salt lake in Crimea, Ukraine in 2007, and deposited in the St. Petersburg Culture Collection (CALU) under the accession number 1533 (Hrouzek et al., 2016). Later, in 2015, it was deposited by P. Hrouzek in the Culture Collection of Algae and Protozoa (CCAP) under the accession number 1453/38. The strain was tested for cytotoxic effects against mammalian cells, including tumor cells (Hrouzek et al., 2016; Voráčová et al., 2017; Chmelík et al., 2019).

Cultivation of *Nostoc* was performed in 250 mL Erlenmeyer flasks placed on a multi-color cultivation bench described previously (Bernát et al., 2021). The flasks were shaken daily to prevent excessive sedimentation of the cells. All cultivations were performed in ambient air at 24 °C in BG-11 medium (Rippka et al., 1979) in a batch regime. Inoculum cultures were cultivated under continuous cool white fluorescent lamps (25 μmol photons m^−2^ s^−1^). Prior to cultivation under narrow-band LEDs, all cultures were diluted with fresh BG-11 medium to an optical density of OD_750_ = 0.2 (determined using a Specord 210 Plus spectrophotometer, Analytik Jena, Germany).

During the cultivation experiments, illumination was provided in a 12:12 light-dark regime by a custom-built illumination setup equipped with eight distinct types of narrow-band LEDs (spectral profiles shown in Fig. 1A): FD-34UV-Y1 (peak at 435 nm), FD-3B-Y1 (465 nm), FD-32G-Y1 (495 nm), FD-3G-Y1 (520 nm), B08QCMC3K1 (555 nm), FD-3R-Y1 (633 nm), FD-333R-Y1 (663 nm), and FD-34R-Y1 (687 nm). Except for the 555 nm diode, which was manufactured by Nagulagu Co., Ltd. (Shenzhen, China), all light sources were obtained from Shenzhen Fedy Technology Co. Ltd. (Shenzhen, China). The PAR from each LED was adjusted to photon flux density (PFD) of 25 μmol photons m^-2^ s^-1^.

**Fig. 1.**
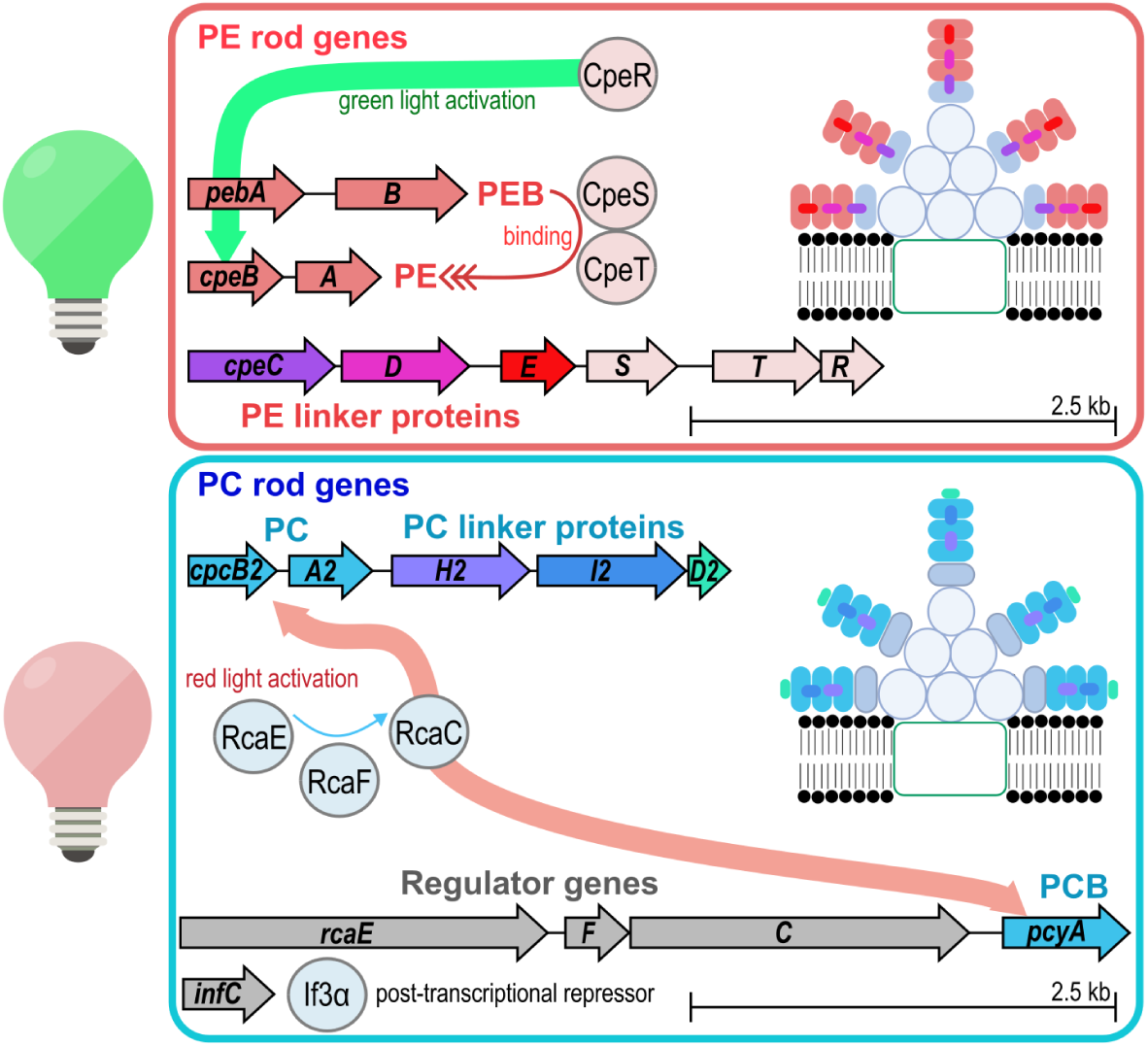
Complete genetic machinery for CA3 in *Nostoc*. The upper panel shows homologs of the genes for phycoerythrin subunits (PE), phycoerythrobilin (PEB) and associated linker proteins, PE maturation proteins (CpeS, CpeT), and the green light-inducible transcriptional activator of PE rod proteins (CpeR) identified in the *Nostoc* genome. The lower panel shows homologs of the genes for phycocyanin subunits (PC), phycocyanobilin (PCB), and associated linker proteins, as well as the red-light-inducible regulatory cascade (RcaEFC) activating the transcription of PC rod genes, and the post-transcriptional repressor (*infC*, encoding IF3α). The red and blue contours in the upper and lower panels match the color appearance of *Nostoc* cells under green and red light, respectively.

### 2.2 Growth rate, cell composition

The specific growth rate was determined by fitting the OD_750_ values (measured using a Specord 210 Plus spectrophotometer) with an exponential regression model. Phycobilisomes were quantified according to a previously described protocol (Zavřel et al., 2018a). Chl *a* and carotenoids were extracted from cellular pellets with acetone and quantified using a Shimadzu Prominence HPLC system (Shimadzu, Japan), following a method described previously (Bernát et al., 2021; Zavřel et al., 2024). The HPLC system was equipped with a Phenomenex Synergi Hydro-RP 250 × 4.6 mm column with a particle size and a pore size of 4 µm and 80 Å, respectively. Elution was performed using a linear gradient from solvent A (acetonitrile:water:trimethylamine, 9:1:0.01) to solvent B (ethyl acetate) for 25 min at a flow rate of 1 mL min^−1^ at 25°C. The relative abundance of proteins, lipids, and carbohydrates was estimated by Fourier-transform infrared spectroscopy (FTIR) of the lyophilized cellular pellets, using a Nicolet iS10 spectrometer (Thermo Fisher Scientific), following previously described protocols (Felcmanová et al., 2017; Rabouille et al., 2021). In brief, the amplitudes of the ∼1735 cm^-1^, ∼1652 cm^-1^, and ∼1152 cm^-1^ FTIR bands were evaluated to determine the content of lipids, proteins, and carbohydrates, respectively. Since the variability of cellular protein content between samples was identified as the lowest among the tested macromolecular pools (data not shown), all spectra were normalized to the protein band at ∼1652 cm^-1^. The length of the fatty acid chains was estimated from the ratio of the amplitudes of asymmetric vibrations of CH_3_ at ∼2960 cm^-1^ and CH_2_ at ∼2925 cm^-1^.

### 2.3 Absorption and fluorescence spectroscopy

Whole-cell absorption spectra were recorded using a Specord 210 Plus spectrophotometer (Analytik Jena, Jena, Germany). To correct for light scattering by the cellular matter, four slices of tracing paper were placed in front of both the sample and reference cuvettes. To calculate the photosynthetically usable radiation (PUR) for each narrow-band LED, the baseline-corrected absorbance spectra were used:

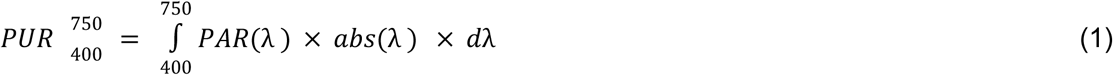

where *PAR(λ)* represents the irradiance of the cultivation LEDs, and *abs*(*λ)* represents absorbance spectra of the corresponding *Nostoc* cultures at wavelength λ, and dλ is the wavelength differential across the 400–750 nm range.

The 3D fluorescence excitation-emission maps were recorded at 77 K using a Jasco FP-8550 spectrofluorometer (Jasco, Tokyo, Japan). 10 mL culture aliquots were filtered through GF/B filters (Whatman, Maidstone, UK), flash-frozen in liquid nitrogen, and stored at −80 °C. Right before the measurements, a slice from each filter (∼1 cm × 0.3 cm) was cut to fit the metal sample holder, which was then inserted into the transparent finger of the Dewar flask. Fluorescence maps were recorded over the excitation and emission ranges of 350–650 nm (step: 5 nm) and 500–800 nm (step: 0.5 nm), respectively. The obtained fluorescence maps allowed us to distinguish, on a relative basis, Chl *a* fluorescence originating either from PSII (Chl-PSII) or PSI (Chl-PSI), the fluorescence of PBS coupled to either PSII (PBS-PSII) or PSI (PBS-PSI), or functionally uncoupled from both photosystems (PBS-free), as well as the ratio of phycocyanin (PC) to phycoerythrin (PE) within the PBS, according to the following equations:

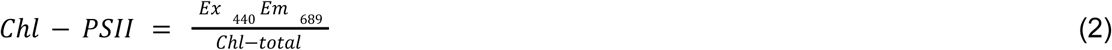

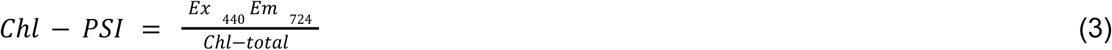

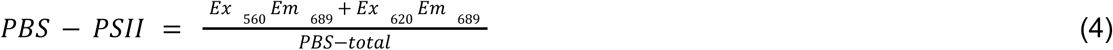

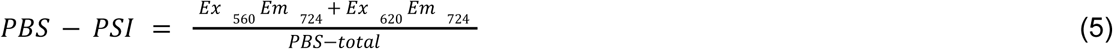

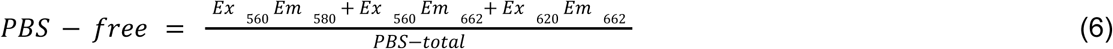

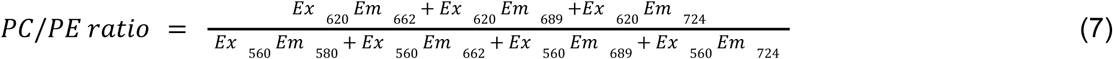

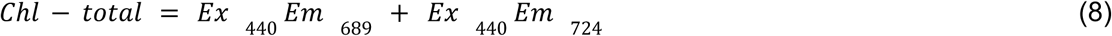

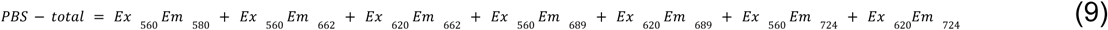

where Ex and Em represent excitation and emission, respectively, with the subscripts representing the corresponding wavelengths. For further details, see Fig. 4.

### 2.4 PSII activity measurement

The activity of PSII was probed using a Multi-Color PAM (Walz, Effeltrich, Germany) by recording the functional absorption cross-section of PSII (σ_II_; (Schreiber et al., 2012)), slow Chl *a* fluorescence kinetics designed to estimate the PSII-mediated electron transport rate (ETR), state transition (ST) and NPQ, as well as fast Chl *a* fluorescence induction (OJIP transients).

σ_II_ was measured using the default Multi-Color PAM script *Sigma1000cyano*, the trigger file *Sigma1000*, and the fitting protocol described in detail previously (Schreiber et al., 2012) in cultures with a Chl *a* concentration of 700 ± 300 μg L^-1^. The σ_II_ values were used to calculate the electron transport rate based on the quantum absorption and yield of PSII, according to the following equations:

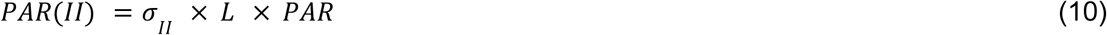

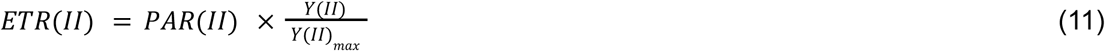

where PAR is the PAR intensity, PAR(II) is the rate of PAR absorption by PSII, L is Avogadro’s constant, ETR(II) is the rate of PSII-mediated electron transport (units: e^-^ PSII^−1^ s^−1^), Y(II) is the effective PSII quantum yield under AL, and Y(II)_max_ is the maximum PSII quantum yield in the dark-acclimated state, under which σ_II_ was determined (Schreiber et al., 2012). The Y(II) values, used for the ETR(II) calculation, were determined during the slow Chl *a* fluorescence kinetics measurements, as follows:

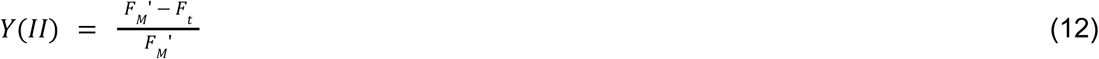

where F_M_’ and F_T_ are the maximum and steady-state fluorescence yields under actinic light (AL), respectively. Y(II) and ETR(II) were determined in cultures acclimated to 625 nm AL, i.e. in State 2. To induce ST, the cell suspension was illuminated either by a 480 nm AL at a PFD of 80 μmol photons m^−2^ s^−1^ for 3 min (inducing State 1) or by a 625 nm AL at a PFD of 50 μmol photons m^−2^ s^−1^ for 5 min (inducing State 2; (Calzadilla and Kirilovsky, 2020)). The ST kinetics were assessed by fitting the dynamics of the F_M_’ values following the change in AL quality with an exponential model using the least-squares fitting method:

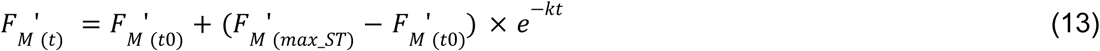

where F_M_’_(t0)_, F_M_’_(t)_, and F_M_’_(max_ST)_ are initial, actual, and maximum fluorescence yields induced by SP under AL upon AL wavelength shift, *t* is time under AL and *k* is the rate constant for ST (units: s^-1^). To induce NPQ, 480 nm light at a PFD of 1 800 μmol photons m^−2^ s^−1^ was additionally applied for 2 min. NPQ was calculated from the maximum fluorescence yield in steady-state under high AL (F_M_’) and during the entire course of each measurement (F_M_’_(max)_; (Bernát et al., 2018), typically recorded at the onset of 625 nm AL) as:

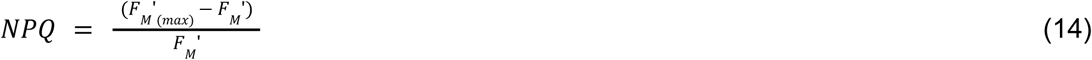

The OJIP curves were recorded using a Multi-Color PAM after 15 min of dark acclimation, with a 625 nm measuring light (ML) and a saturation pulse (SP) at an intensity of 2 200 μmol photons m^−2^ s^−1^. From the fluorescence induction traces, the following parameters were derived:

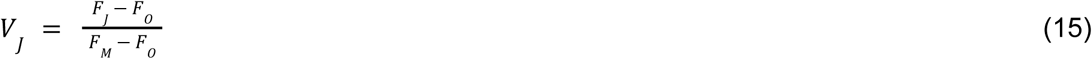

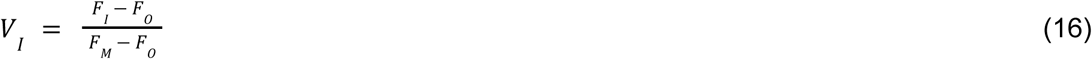

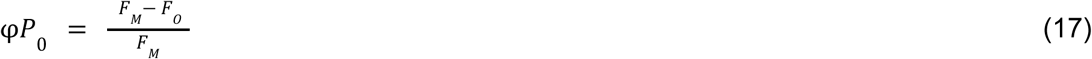

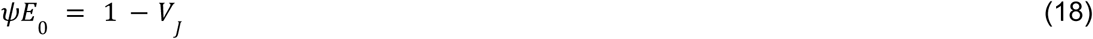

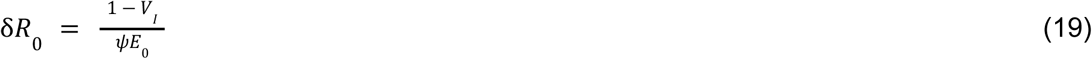

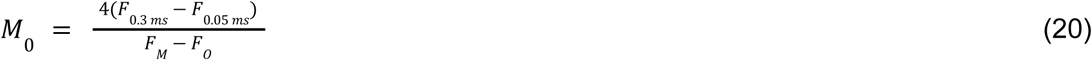

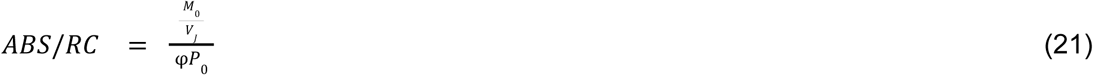

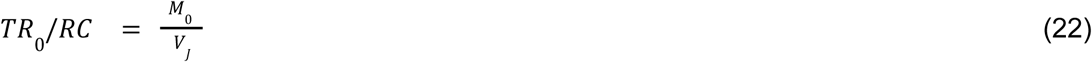

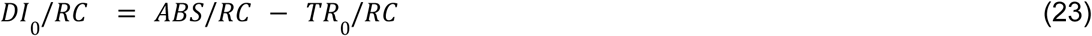

where F_O_, F_J_, F_I_ and F_M_ refer to the fluorescence yields at the O, J and I steps of the OJIP curve, and the maximum fluorescence yield, while V_J_ and V_I_ represent the relative variable fluorescence yields at the J and I steps, respectively. The parameters φP_0_, ψE_0_ and δR_0_ represent the maximum quantum yield of primary photochemistry and the efficiencies with which a PSII-trapped electron is transferred from quinone A (Q_A_ ^−^) to PQ and from plastoquinol (PQH_2_) to the final electron acceptors, respectively. The parameter M_0_ represents the initial slope of the OJIP curve, while the parameters ABS/RC, TR_0_/RC and DI_0_/RC represent the apparent antenna size of an active PSII, the maximum trapped exciton flux per active PSII, and the flux of energy dissipated in processes other than exciton trapping per active PSII, respectively (Stirbet et al., 2018).

### 2.5 PSI activity measurement

PSI kinetics were probed using a Dual-PAM-100 fluorometer (Walz, Effeltrich, Germany). Culture aliquots were filtered through glass fiber filters (GF/B, Whatman). Wet filters were placed between two microscope glass slides embedded in a DUAL-B leaf holder (Zavřel et al., 2024) and dark-acclimated for 10-15 min prior to the measurements. PSI limitation on both the donor and acceptor sides was probed in the SP-analysis mode (using the Fluo+P700 measuring mode). The rate of electron transport through PSI was determined in the FastAcquisition mode in the absence of inhibitors to estimate the total electron flow through PSI, and in the presence of 10 μM DCMU to estimate the rate of cyclic electron flow around PSI (CEF-PSI, units: s^-1^) by inhibiting electron flow through PSII. CEF-PSI expresses the rate of P_700_ ^+^ re-reduction kinetics, obtained by a single-exponential decay fitting of the 830 nm absorption signal, as measured in the dark after oxidizing PSI with a red SP (635 nm, 100 ms; (Zavřel et al., 2018b)).

### 2.6 Confocal microscopy

Confocal images of *Nostoc* cells were acquired using a TCS SP8 DMI confocal laser scanning microscope (Leica Microsystems Inc., Wetzlar, Germany), equipped with a triple-band 488/552/638 main beam splitter to separate the reflected excitation light from the fluorescence emission. Images with a resolution of 1024 × 1024 pixels, covering an area of 123.02 μm × 123.02 μm, were collected using an ‘HC PL APO CS’ 63× oil immersion objective (numerical aperture, NA: 1.4) with a 1.5× zoom factor. Prior to imaging, 1 mL culture aliquots were sampled from the Erlenmeyer flasks and centrifuged (2500 × *g*, 5 min, 25°C). The supernatant was partially discarded and 20 µL of the concentrated sample was placed onto a microscope slide covered with a coverslip, and mounted inverted on the microscope stage. Chl *a* and PE autofluorescence were excited using the 488 nm laser line and detected over the 690–790 nm and 550–610 nm spectral windows, respectively. The PBS autofluorescence was excited using the 638 nm laser line and detected over the 650–680 nm spectral window. The sensitivity of the detectors (photomultiplier tubes, PMTs) was kept constant for all measurements: the PMT1 (PE) gain was set to 650 V, whereas the PMT2 (Chl *a*) and PMT3 (PBS) gains were set to 625 V and 600 V, respectively. The scanning speed was set to 400 Hz, and the pinhole size was maintained at 0.80 AU throughout all measurements. The distance between consecutive focal planes was 0.2 µm during optical sectioning for Z-stack acquisition.

### 2.7 Bioinformatics analysis

Previously published raw genome sequencing data of *Nostoc* (NCBI BioProject accession no. PRJNA266493, Sequence Read Archive accession no. SRX752833) were assembled using CLC Bio Genomics Workbench v. 7.5 as described previously (Voráčová et al., 2017) and deposited in the NCBI GenBank database under accession no. JAWJTF000000000. The whole genome draft was explored for the presence of the Orange Carotenoid Protein (OCP), and the genes encoding protein building blocks and machineries typical for individual CA types found in cyanobacteria (Sanfilippo et al., 2019): BolA, CcaR, CcaS, CpcA2, CpcB2, CpcC, CpcD2, CpcI2, CpcL, CpeA, CpeB, CpeC, CpeD, CpeE, CpeR, CpeS, CpeT, CphA, DpxA, IflA, InfC, MpeY, MpeZ, MreB, PebA, PebB, PecA, RcaC, RcaE, and RcaF. Genomic loci corresponding to these proteins were identified using *tblastn* (with an *E-*value cut-off of 1 × 10^-10^) against a custom BLAST database built from the *Nostoc* genome assembly in Geneious Prime v. 2026.0.2 (www.geneious.com), using a selection of functionally annotated cyanobacterial proteins as queries. These queries were collected from the NCBI Protein database entries of a set of previously studied cyanobacterial species listed by (Hirose et al., 2019), e.g., *Anabaena* sp. PCC 7120, *Fremyella diplosiphon* Fd33/PCC 7601, *Nostoc flagelliforme* CCNUN1, *Nostoc punctiforme* PCC 73102, *Synechococcus* sp. A15-24, and *Synechocystis* sp. PCC 6803. Genomic regions of *Nostoc* containing BLAST search hits were annotated using Glimmer3 (Delcher et al., 2007); ORFs were translated and analyzed using *blastp* and the Conserved Domain Database search against NCBI to verify their identity. The complete set of genes corresponding to the CA3 type and OCP was mapped onto *Nostoc* genomic scaffolds in Geneious Prime and visualized using Inkscape v. 1.3 (www.inkscape.org).

### 2.8 Statistical analysis

Statistical analyses were conducted according to the workflow described previously (Zavřel et al., 2024). All tests were performed using R Statistical Software (R Core Team, 2022). When both the assumptions of normality and homogeneity of variances were satisfied (Fox and Weisberg, 2011), ANOVA followed by Tukey’s HSD post hoc test was used. If only homogeneity of variances was met, the Kruskal–Wallis test with multiple comparisons (Giraudoux et al., 2023) was used. When only data normality was satisfied, Welch’s one-way ANOVA was used, followed by pairwise t-tests with Benjamini–Hochberg correction. The number of replicates was 3–4 for all growth lights throughout all experiments except for the parameters PSI-CEF (n = 9-12), growth rate (n = 4-5), and cell diameter (n = 93-164). The significance level was set to *p* < 0.05.

## 3. Results

### 3.1 *Nostoc* sp. CCAP 1453/38 is a CA3 type cyanobacterium

In this work, the light quality acclimation of the cyanobacterium *Nostoc* sp. CCAP 1453/38 was evaluated. *Nostoc* was identified as a chromatic acclimation type 3 (CA3) cyanobacterium, harbouring all essential components of the CA3 genetic machinery in its genome (Fig. 1). CA3-type cyanobacteria (Sanfilippo et al., 2019) are expected to contain the photoreceptor gene *rcaE* that responds to both red and green light and regulates a phosphorelay cascade involving two response regulators: *rcaF* (a single-domain response regulator) and *rcaC* (a large, complex response regulator with multiple domains including DNA-binding capability). CA3 cyanobacteria further possess the *cpeBA* operon encoding α- and β-subunits of phycoerythrin, the *cpeCDESTR* operon encoding phycoerythrin linker proteins and regulatory components, and the *pebBA* operon encoding phycoerythrobilin synthase for chromophore biosynthesis. Expression of these operons is regulated by CpeR, which acts as an activator essential for phycoerythrin gene expression. For phycocyanin production, CA3 cyanobacteria contain the *cpcB2A2* operon encoding specialized phycocyanin subunits preferentially expressed under red light, and the associated linker genes (cpcH2, I2, and D2).

### 3.2 PE overexpression provides fitness advantage over PC-rich strains under green light

To characterize *Nostoc* growth under 400–750 nm wavelength range, the PFD of PAR of all narrow-band LEDs was set to 25 μmol photons m^-2^ s^-1^ (Fig. 2A). As the absorption of the *Nostoc* cells changed within the studied wavelength range (Fig. 2B), the resulting photosynthetically usable radiation (PUR, Eq. (1)) was the highest under violet and green light (Fig. 2C). However, the specific growth rate was the highest under green, red, and near far-red light (555 - 687 nm), whereas it was significantly reduced under blue and violet light (Fig. 2D). The electron transport rate at PSII, ETR(II), was upregulated under green and red light (520–663 nm; Fig. 2E), similar to CEF-PSI which was, however, also upregulated under near far-red light (Fig. 2F). The PSI limitation at the donor side (Y(ND)) was the highest under blue light (Fig. 2G), while the PSI limitation at the acceptor side (Y(NA)) was the highest under green light (Fig. 2H).

**Fig. 2.**
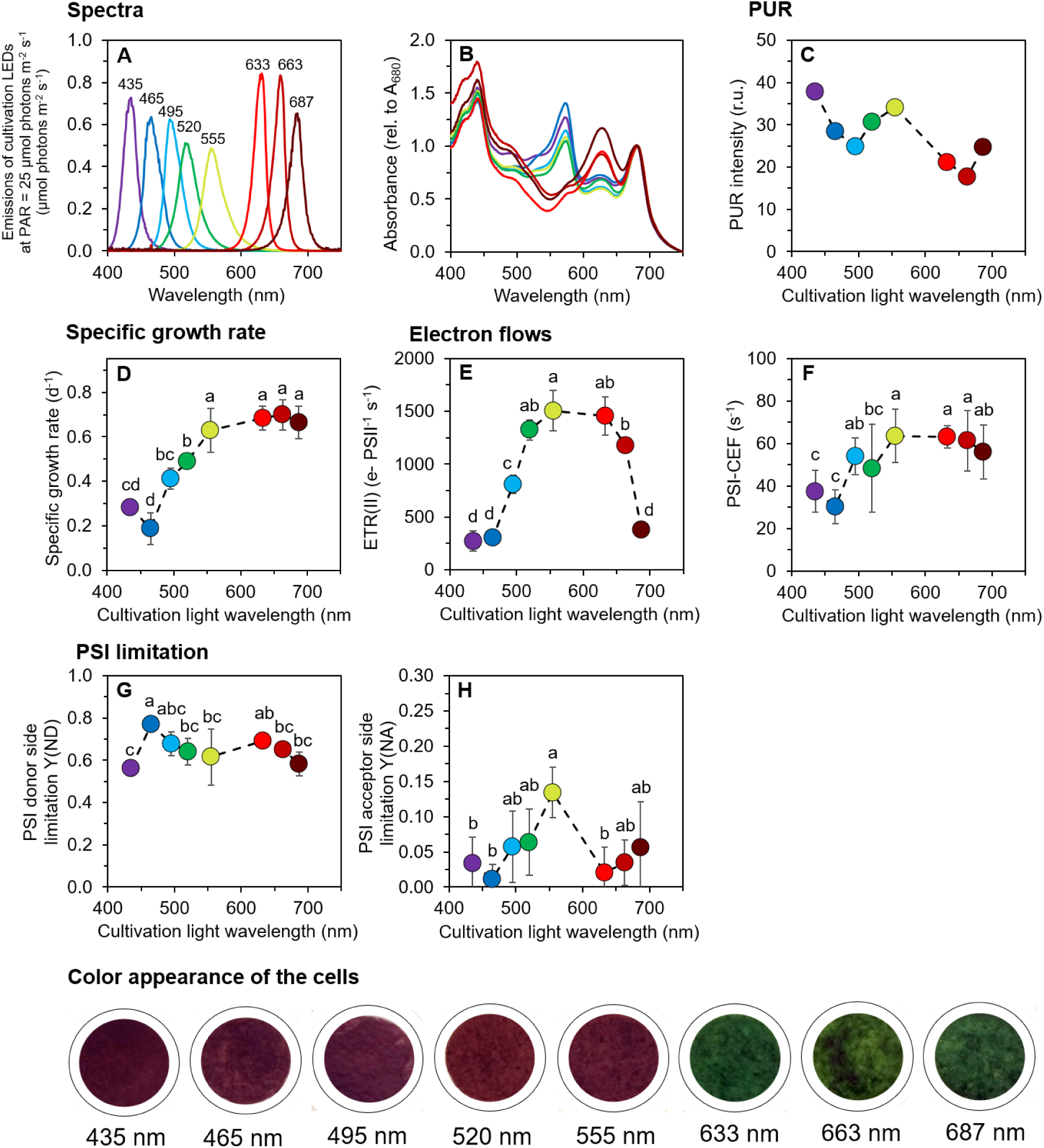
Emission spectra of cultivation LEDs at photosynthetically active radiation (PAR) of 25 μmol photons m^-2^ s^-1^ (A), baseline-corrected absorption spectra of *Nostoc* cultures (B), photosynthetically usable radiation, PUR (panel C; see Eq. (1) for details), specific growth rate (D), electron transport rate through PSII (ETR(II); panel E), cyclic electron flow around PSI (CEF-PSI; panel F), and donor-side (Y(ND); panel G) and acceptor-side limitation of PSI (Y(NA); panel H) in *Nostoc* cells cultivated under narrow-band LEDs. CEF-PSI was determined from P ^+^ _700_ re-reduction kinetics, as measured after saturation pulse (635 nm, 100 ms; see Materials and Methods for details). Values in panels B and D–H represent mean±SD (n = 3–12). Values in panels B and C are shown without error bars for clarity. Different letters above the symbols in panels D–H indicate statistically significant differences within each parameter (p < 0.05).

These results show that the absolute PUR intensity was not the only driver of *Nostoc* growth across the PAR spectrum, and that a relatively high growth rate can be maintained not only through linear electron flow but also *via* CEF-PSI, the intensity of which varied with the preferential excitation of either PSII or PSI (see below). Notably, although PUR was similar under violet (435 nm) and green (520 nm) light, the resulting growth rates differed substantially (Fig. 2D). This discrepancy between absorbed and biologically usable energy points to wavelength-specific differences in how absorbed photons are distributed between PSII and PSI - a recurring pattern across our dataset that is addressed in detail in the Discussion.

### 3.3 Trends in pigmentation and cellular composition in *Nostoc* across the PAR spectrum reveal both unique and shared responses compared with PC- and PE-rich cyanobacteria

The cellular content of all pigments – specifically total PBS, individual phycobiliproteins, Chl *a*, and carotenoids – was upregulated under violet–blue light (435–495 nm) and downregulated under red light (633–687 nm; Fig 3A–C, Supplementary Figs. S1 and S2). These trends were opposite to those observed in the CA1 cyanobacterium *Synechocystis* sp. PCC 6803, which reduced content of all pigments under blue light **(Zavřel et al., 2024)**, as well as in the non-CA strain *Cyanobium* sp. NIVA-CYA 375, which upregulates only PBS but not Chl *a* and carotenoids under violet–blue light **(Kis et al., 2026)**. On the other hand, these trends were similar to those reported for *Cyanobium gracile*, another non-chromatic acclimator **(Bernát et al., 2021)**.

**Fig. 3.**
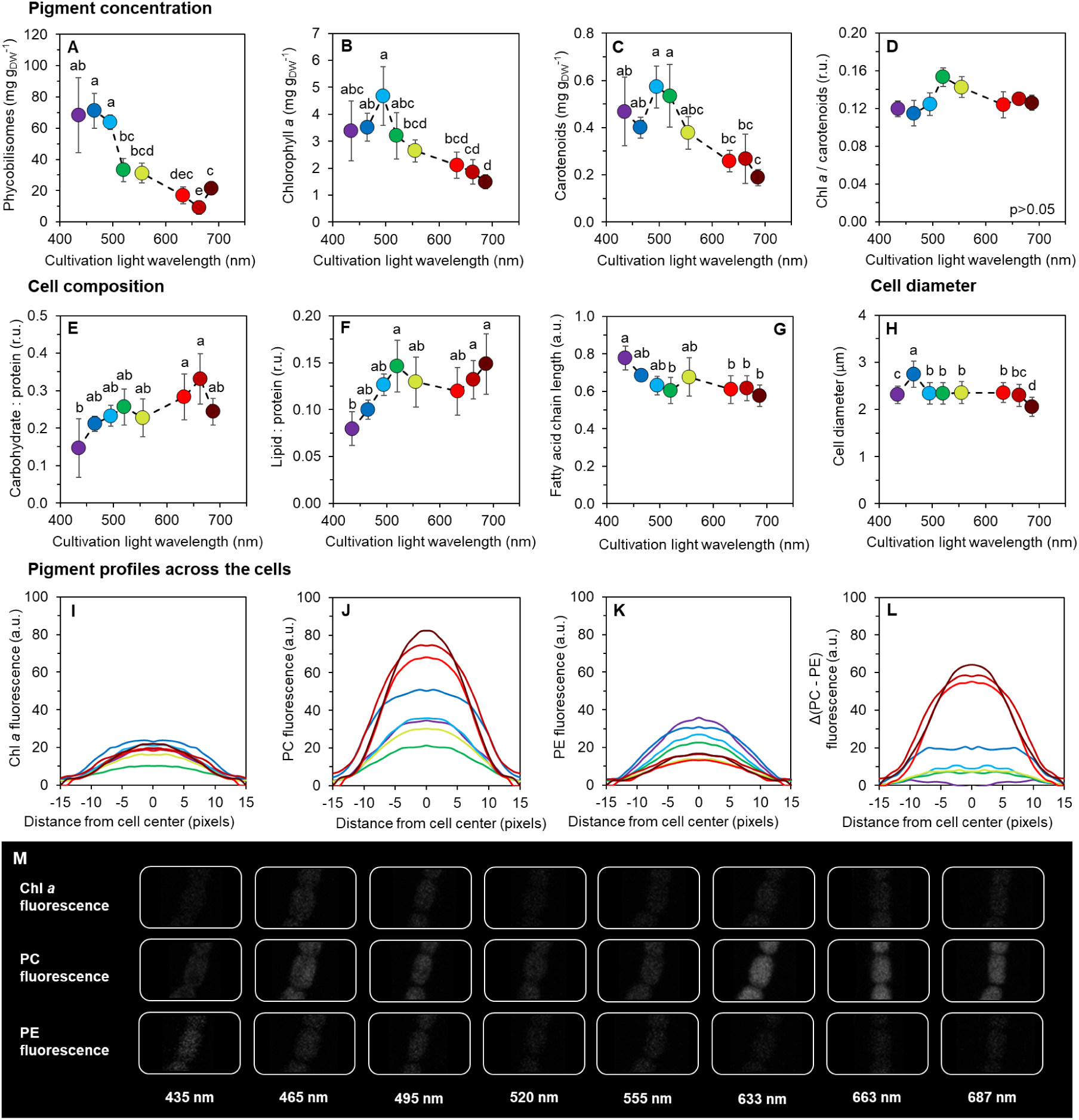
Content and ratio of pigments (A–D) and macromolecular pools (E–G), cell size (H), and pigment profiles (I–M) in *Nostoc* cultivated under narrow-band LEDs. Phycobilisomes (A) and carotenoids (C) were calculated as the sum of phycocyanin, allophycocyanin, and phycoerythrin, and β-carotene, echinenone, and myxoxanthophyll, respectively. Carbohydrate (E) and lipid (F) contents are expressed relative to protein content; the fatty acid chain length (G) was estimated from the ratio of amplitude values of asymmetric CH_3_ and CH_2_ vibration peaks (Supplementary Fig. S4). Analysis of confocal microscopy images allowed for the estimation of cell diameter (H) and pigment profiles across *Nostoc* cells (I–M). All channels in confocal microscopy images were analyzed in a transverse view, i.e., in the plane perpendicular to the filament axis. Values in panels A–H represent mean±SD (n = 3–4 for panels A–G and n = 93–164 for panel H). Values in panels I–L are shown without error bars for clarity (n = 72–151). Different letters above the symbols in panels A–H indicate statistically significant differences within each parameter (p < 0.05). Panels A-C show pigment content relative to the cellular dry weight (DW).

Among the individual carotenoids, *Nostoc* upregulated β-carotene under 435–520 nm light. In contrast, echinenone and myxoxanthophyll were downregulated under 465 nm light. All carotenoids were downregulated under red light (633–687 nm), with the exception of myxoxanthophyll, which was upregulated under 663 nm (Supplementary Fig. S2). The Chl *a*/carotenoids ratio did not change with the wavelength of the cultivation light (Fig. 3D). The PBS/Chl *a* ratio was upregulated under 435–495 nm as well as under 687 nm light (Supplementary Fig. S3); however, under 687 nm light this reflected a sharper reduction in Chl *a* level rather than an increase in PBS content (Fig. 3A).

Light quality affected the macromolecular composition of *Nostoc* cells. Both carbohydrate and lipid contents, expressed relative to protein content, were downregulated under 435 nm light, and the highest concentrations of both macromolecular pools were detected under 633–663 nm light (Fig. 3E, F). In contrast, the relative fatty acid chain length, calculated from the ratio of asymmetric CH_3_ and CH_2_ vibration amplitudes (Supplementary Fig. S4), was downregulated under red light (633–687 nm; Fig. 3G).

Analysis of the confocal microscopy images (bright-field channel) revealed that the diameter of *Nostoc* cells increased under 465 nm light and decreased under violet and near far-red light (435 nm and 687 nm, respectively), compared with other wavelengths (Fig. 3H). Analysis of the confocal fluorescence microscopy images (Fig. 3I–M) allowed for the evaluation of the relative concentration and ratio of Chl *a*, PC, and PE, as well as the intensity of fluorescence signals from these pigments throughout the *Nostoc* cells’ cross-section. The results show that the thylakoid membranes did not form a distinct parietal (ring-like) structure, which is in accordance with the current concept of fascicular thylakoid membrane arrangement typical for *Nostoc* strains (Mareš et al., 2019). The relative amount and ratio of Chl *a*, PC, and PE were in agreement with the trends observed in whole cell absorption spectra (Fig. 2B), absolute pigment content (Fig. 3A, B), and the results of 77K spectrofluorometry measurements (Fig. 4). We note that all channels in the confocal microscopy images, including the bright-field, were analyzed in a transverse view, i.e., in the plane perpendicular to the filament axis.

**Fig. 4.**
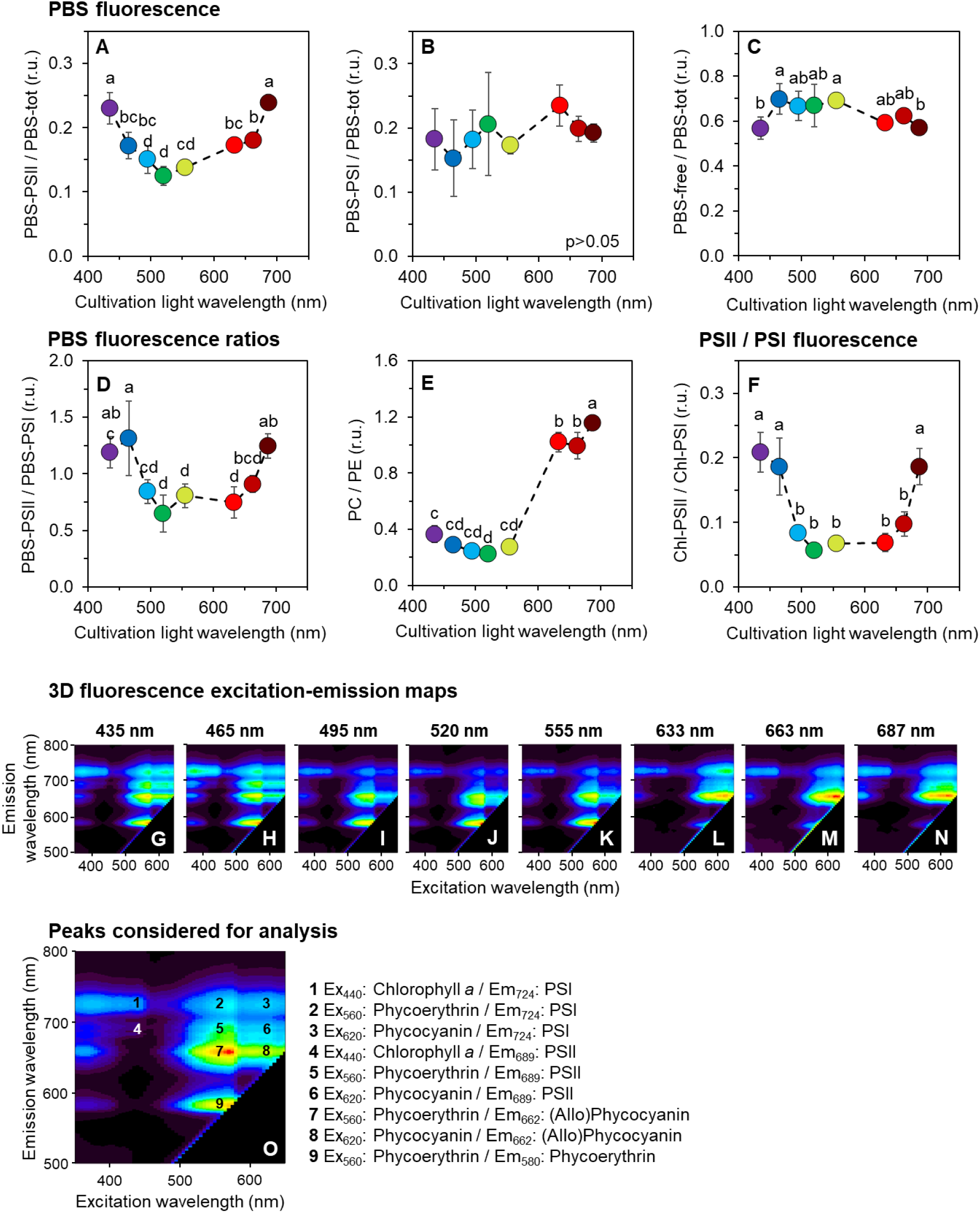
3D fluorescence excitation–emission maps recorded at 77K using *Nostoc* cells cultivated under narrow-band cultivation lights. The derived parameters include fluorescence from phycobilisomes functionally attached to PSII (PBS–PSII; panel A), to PSI (PBS–PSI; panel B), and from phycobilisomes functionally uncoupled from both photosystems (PBS–free; panel C). Further, the ratios of PBS–PSII/PBS–PSI (D), phycocyanin/phycoerythrin (E), and PSII/PSI (F) were calculated. Panels G–N show representative 3D fluorescence excitation–emission maps of cultures grown under all tested cultivation lights; panel O shows the peaks used for 77K spectra analysis (for further details on parameter calculation, see Eqs. (2–9)). Values in panels A–F represent mean±SD (n = 3–4); different letters above the symbols indicate statistically significant differences within each parameter (p < 0.05).

### 3.4 CA3 reduces both the need and options to fine-tune light harvesting under green light

#### 3.4.1 Excitation–emission spectra

To further understand growth limitation across the entire PAR range, 3D fluorescence excitation–emission maps were recorded at 77 K. Analysis of the fluorescence spectra revealed that the functional coupling of PBS to PSII (PBS–PSII) was enhanced under 435 nm and 687 nm light (Fig. 4A), which was accompanied by a reduction in the PBS pool functionally uncoupled from both PSII and PSI (PBS–free; Fig. 4C). Energy transfer from PBS to PSI (PBS–PSI) was independent of the cultivation light wavelength (Fig. 4B). Interestingly, the PBS–PSII/PBS–PSI ratio was upregulated under 435 nm, 687 nm, and also 465 nm light (Fig. 4D), mirroring the trend observed for the PSII/PSI ratio (Fig. 4F). For details on individual PSII and PSI fluorescence changes within the studied wavelength range, see Supplementary Fig. S5; fluorescence excitation and emission spectra are summarized in Supplementary Figs. S6 and S7, respectively.

These results demonstrate the regulation of light harvesting under wavelengths poorly absorbed by the light-harvesting antennas, aiming to alleviate excitonic imbalance and optimize the ATP/NADPH ratio. The upregulation of PBS–PSII under violet–blue and near far-red light has been described previously for CA1 and non-CA strains (Zavřel et al., 2024; Kis et al., 2026). Similarly, PBS–PSII upregulation under green light, compared with red light, has been observed in the CA1 strain *Synechocystis* (Zavřel et al., 2024), pointing to attempts to optimize green light harvesting in PC-rich strains. Since *Nostoc* was able to harvest green wavelengths efficiently *via* PE (Fig. 2B), it was not forced to maintain a relatively high amount of PSII and PBS–PSII under 495–520 nm light, unlike *Synechocystis* sp. PCC 6803 (Zavřel et al., 2024).

The 3D fluorescence analysis further allowed for the evaluation of fluorescence originating from PC and PE individually (Fig. 4O; for details, see Eqs. (2–9)). The resulting PC/PE ratio was upregulated under red light (633–687 nm), in line with spectrophotometry (Supplementary Fig. S1) and confocal microscopy results (Fig. 3J–M).

#### 3.4.2 State transition rates and NPQ

Besides studying the functional coupling of PBS to PSII or PSI, slow Chl *a* fluorescence transients were recorded, allowing for the estimation of the rate and extent of state transitions, as well as the magnitude of NPQ. To induce State 1 and State 2, *Nostoc* cultures were illuminated by weak blue (480 nm) and red (625 nm) AL of the Multi-Color PAM, respectively (Fig. 5H; for AL spectra, see Supplementary Fig. S8). Similar to other cyanobacteria, the State 2→1 rate was faster compared to that of the State 1→2 rate (Calzadilla and Kirilovsky, 2020). Both rates exhibited a specific dependence on the wavelength of the cultivation light (Fig. 5A, B), which differed from previous results in *Synechocystis*, where both rates were the slowest under blue and the fastest under red cultivation light (Zavřel et al., 2024). In *Nostoc*, the State 1→2 rate, determined under 625 nm AL, was the slowest under 435 nm, 465 nm, and 687 nm light and the fastest under 520 nm and 633 nm light (Fig. 5A). The State 2→1 rate, measured under 480 nm AL, showed a partially opposite trend, being the fastest under 435 nm and 465 nm light (Fig. 5B).

**Fig. 5.**
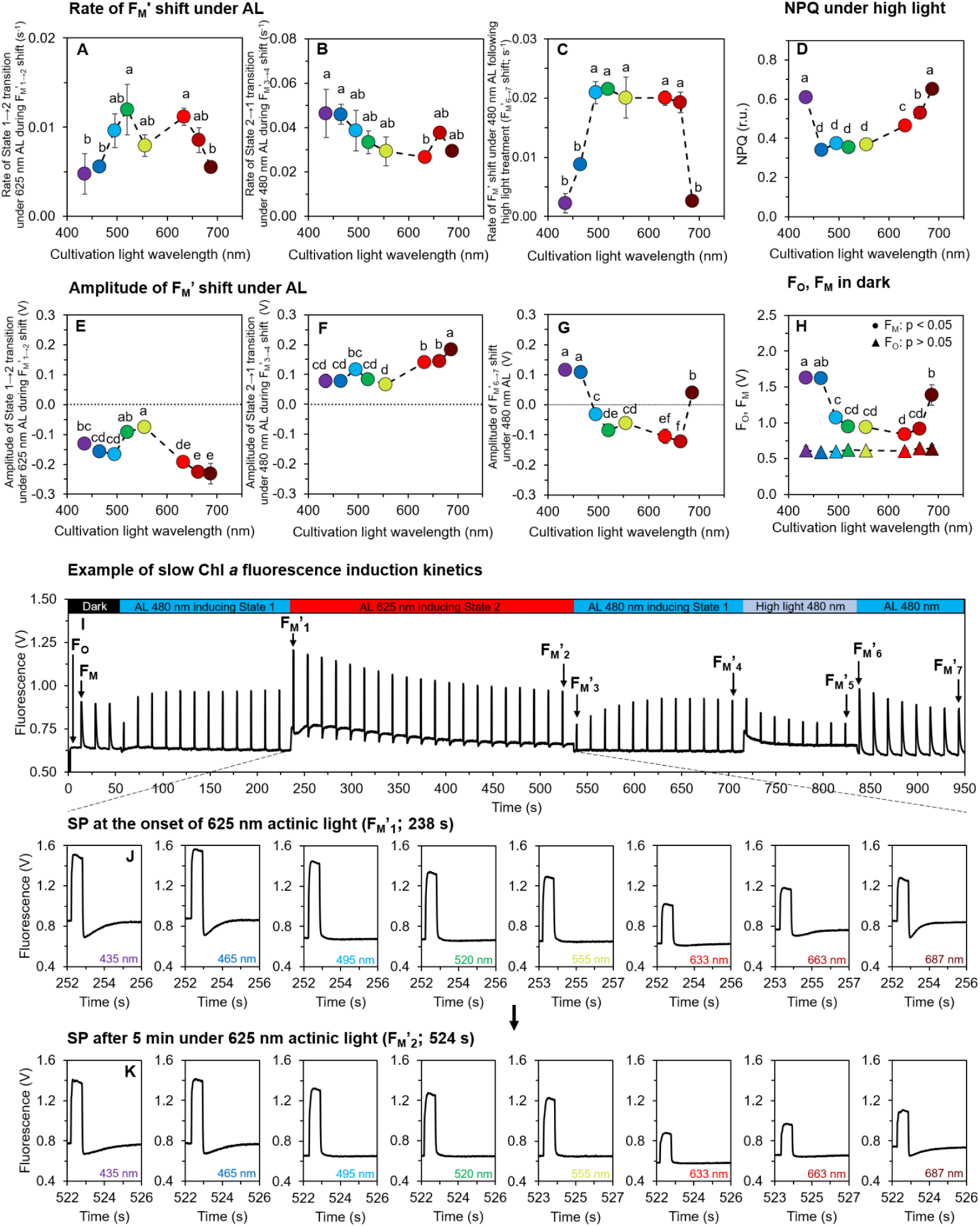
Rates and extent of state transitions, NPQ, F_O_ and F_M_, and fluorescence kinetics after SP application under 625 nm AL in *Nostoc* cells cultivated under 435–687 nm narrow-band LEDs. State 1→2 (A) and State 2→1 transition rates (B) were estimated from the dynamics of F_M_’ under 480 nm and 625 nm AL, respectively (see Eq. (13) for details). PBS (re)coupling to PSII after HL treatment was estimated from the rate of F_M_’ shift (C) after the NPQ measurement under HL (D; according to the measurement protocol shown in panel I). To evaluate the extent of ST under 625 nm and 480 nm AL, differences in F_M_’ values according to the measuring protocol (I) were assessed (E and F, respectively). Similarly, the extent of PBS (re)coupling after the NPQ determination was assessed from the F_M_’ difference subsequent to the HL treatment (G). Panel H shows F_O_ and F_M_ values, measured after 15 min of dark acclimation. Values in panels A–H represent mean±SD (n = 3–4); different letters above the symbols indicate statistically significant differences within each parameter (p < 0.05). Panel I shows a representative Chl *a* fluorescence trace from which the ST rates and NPQ were calculated. Prior to each measurement, *Nostoc* cultures were dark-acclimated for 15 min. Fluorescence traces from all cultivation light conditions are summarized in Supplementary Fig. S10, and additional parameters are presented in Supplementary Fig. S11. Emission spectra of the 480 nm and 625 nm LEDs of the Multi-Color PAM are summarized in Supplementary Fig. S8. We note that the amplitude of F_M_’ under 480 nm AL is underestimated in MC-PAM (Pfennig et al., 2024; Zavřel et al., 2024).

State 2 is characterized by an increased absorption cross-section of PSI (σ_I_). The relatively slow State 1→2 transition in *Nostoc* cultures cultivated under violet and blue light (Fig. 5A) was thus likely related to the elevated PSII/PSI ratio under these wavelengths (Fig. 4F, Supplementary Fig. S5). In contrast, State 1 is characterized by an increased absorption cross-section of PSII (σ_II_). The relatively fast State 2→1 transition under violet and blue light can be analogously attributed to this elevated PSII pool relative to PSI (Fig. 5F, Supplementary Fig. S5). Under 687 nm light, both ST rates were low. This can be explained by the combination of low PSI levels (Supplementary Fig. S5) and low PBS content (Fig. 3A).

Furthermore, ST exhibited similar rates under green (520–555 nm) and red (633–663 nm) cultivation light (Fig. 5A,B), suggesting a reduced need and/or reduced options for light-harvesting optimization under the combination of low PSII levels (Supplementary Fig. S5) and low PBS content (Supplementary Fig. S3). However, the absolute shifts in F_M_’ values during ST were lower under green light, compared with red light (Fig. 5E, F). This was likely related to the relatively high PC/PE ratio in PBS under red cultivation light (Figs. 2B, 3L, and 4E): PC-rich PBS were able to capture the 625 nm AL with higher efficiency than PE-rich PBS, which can be expected to result in a more intense ST. These results demonstrate that the fluorescence traces need to be interpreted with respect to PBS composition and the specific AL wavelength range used.

In addition to the ST rates, the fluorescence transients allowed for the evaluation of the magnitude of NPQ. In cyanobacteria, NPQ is typically mediated by the orange carotenoid protein (OCP) or its homologs (Kerfeld et al., 2017), the sequence of which was also identified in the genome of *Nostoc* (Supplementary Fig. S9). OCP-mediated NPQ was triggered by blue AL of high intensity (Fig. 5H; Supplementary Fig. S10). The highest NPQ levels were found in cultures cultivated under 435 nm and 687 nm cultivation light (Fig. 5D), which is again likely a result of high PSII content, relative to PSI (Fig. 4F, Supplementary Fig. S5); with more PSII, more excitation energy is transferred from PBS to PSII, thereby leading to a higher NPQ. Slightly lower NPQ levels were found in cultures cultivated under 633 nm and 663 nm light (Fig. 5D). Under these conditions, σ_II_ at the onset of high-light (HL) treatment was estimated to be relatively high, as inferred from the highest absolute shifts in F_M_’ values during the State 2→1 transition preceding the HL treatment (Fig. 5F, Supplementary Fig. S10).

Interestingly, cultures cultivated under 465 nm light exhibited lower NPQ levels than those grown under 435 nm and 687 nm lights (Fig. 5D), despite a similar PSII content and PSII/PSI ratio (Supplementary Fig. S5). This could have several origins, including differences in the expression levels of OCP or the CBB cycle enzymes, respiratory electron flow, CEF–PSI, or the ability to uncouple PBS from PSII. Under 465 nm light, echinenone, one of the structural components of OCP (Kerfeld et al., 2017), was downregulated in *Nostoc* (Supplementary Fig. S2). The increased F_t_ values under HL in the 465 nm culture (Supplementary Fig. S10) suggest that PBS were also uncoupled to a greater extent than in the other cultures (Remelli and Santabarbara, 2018). Furthermore, the highest Y(ND) and the lowest Y(NA) levels were observed in the 465 nm culture (Fig. 2G, H), suggesting potential additional differences downstream of PSII. Taken together, these results suggest that the relatively lower NPQ in the 465 nm culture, compared with the 435 nm and 687 nm cultures, likely has multiple origins.

After the NPQ determination under HL, the PFD was lowered to 80 μmol photons m^-2^ s^-1^. The kinetics of the F_M_’ shift were the slowest in cultures cultivated under 435 nm, 465 nm, and 687 nm light (Fig. 5C). Unlike the State 2→1 transition (Fig. 5B), this F_M_’ shift reflected not only PBS (re)coupling to PSII (expected to be strongest under 465 nm light; (Supplementary Fig. S10) but also relaxation of the OCP-NPQ (highest under 435 nm and 687 nm lights; Fig. 5D). Interestingly, under 435 nm, 465 nm, and 687 nm cultivation light, F_M_’ levels increased after the HL treatment, whereas an opposite trend was observed under all other cultivation wavelengths (Fig. 5G). The kinetics of F_M_’ recovery after the HL treatment are typically measured in darkness and comprise several components: (1) an energy-dependent component related to the built-up of a pH gradient under HL; (2) a photoinhibitory component related to PSII damage under HL; (3) a ST component, which can include rapid PBS detachment from PSII in cyanobacteria (Zavřel et al., 2026); and (4) the relaxation of heat dissipation, including zeaxanthin epoxidation in plants and algae (van Amerongen and Croce, 2025) and OCP relaxation in cyanobacteria (Liu et al., 2018). In our experiments, blue AL was applied after the HL treatment, and the true F_M_ value was not estimated, e.g. *via* DCMU treatment (Stirbet et al., 2019). Therefore, individual NPQ components were not quantified. Nevertheless, the F_M_’ decline after HL in the 495–663 nm cultures can most likely be attributed to ST. As there were generally fewer PSII than PSI centers in *Nostoc* cells (Fig. 4F), the blue AL, captured predominantly by Chl *a*, primarily excited PSI, which is expected to induce State 1 *via* σ_II_ increase. However, σ_II_ was likely reduced compared with the HL treatment (Supplementary Fig. S10). This may represent an advantageous photoprotective strategy for PSII during the recovery phase. The σ_II_ increase in cultures grown under 435 nm, 465 nm, and 687 nm cultivation lights suggests either reduced options for PBS uncoupling from PSII (due to higher PSII content) or a higher involvement of energy-dependent and photoinhibitory NPQ components under HL, the relaxation of which allows for higher fluorescence yields.

In addition to ST and NPQ, a transient drop in fluorescence below the steady-state level was observed at the onset of the 625 nm AL, inducing State 1→2 transition (Fig. 5J). This drop was most pronounced in cultures cultivated under 435 nm, 465 nm, and 687 nm light, and it was partially diminished after 5 min under the 625 nm AL (Fig. 5K). Such temporal decline was previously assigned to a temporal limitation of linear electron flow at the PSII acceptor site (Tsuyama et al., 2004). This drop can thus be related to a relatively low PSI content, consistent with observations in *Synechocystis* sp. PCC 6803 and *Cyanobium* sp. NIVA-CYA 375 (Zavřel et al., 2024; Kis et al., 2026).

#### 3.4.3 Shifting PSII/PSI and PC/PE ratios require careful interpretation of OJIP transients

To further understand photosynthetic efficiency across the studied wavelength range, fast Chl *a* fluorescence induction transients – the so-called OJIP curves – were recorded. The OJIP curves, and especially the JIP-test can provide semi-quantitative information about many parts of PETC (Stirbet et al., 2018). OJIP transients can be measured in both dark- and light-acclimated states (Zavřel et al., 2026). Here, the OJIP curves were recorded only in the dark-acclimated state (Fig. 6A), since pre-illumination by the specific cultivation wavelengths can be expected to affect the internal redox state of PSII differently (i.e., the Q_A_/Q_A_^−^ ratio). In such cases, the capacity of fluorescence transients to reflect PETC parameters is substantially reduced (Zavřel et al., 2026).

**Fig. 6.**
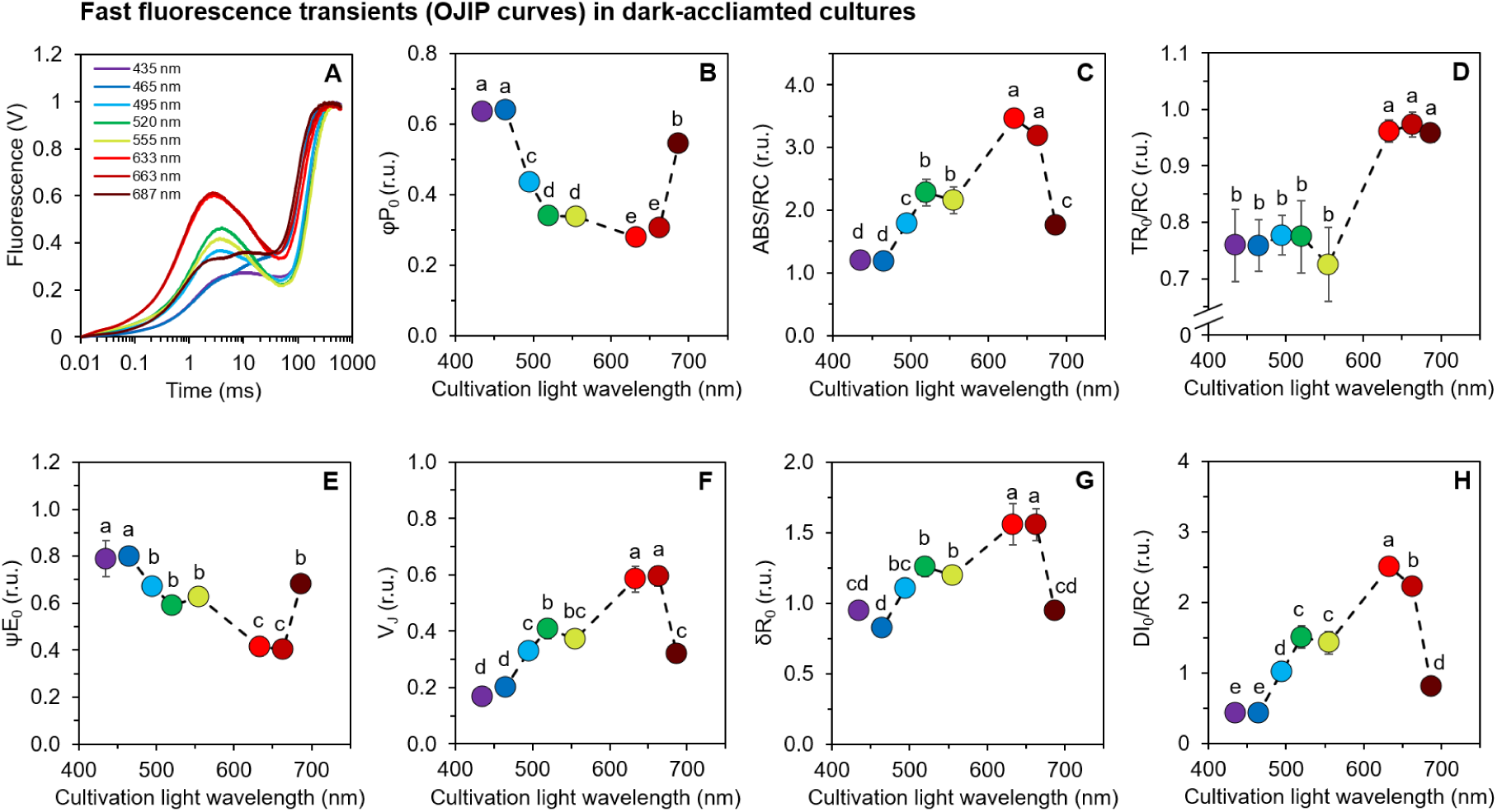
Fast Chl *a* fluorescence induction kinetics (OJIP curves; A) measured in *Nostoc* cultures cultivated under narrow-band LEDs. The derived parameters represent maximum quantum yield of primary PSII photochemistry (φP_0_; B), the apparent antenna size of an active PSII (ABS/RC; C), the maximum trapped exciton flux per active PSII (TR_0_/RC; D), the efficiency with which a PSII-trapped electron is transferred from Q ^-^ to PQ (ψE ; E), the relative variable fluorescence at the J-step, related to the redox state of the PQ pool (V_J_; F), the efficiency with which an electron from PQH_2_ is transferred to final PSI acceptors (δR_0_; G), and the flux of energy dissipated in processes other than trapping per active PSII (DI_0_/RC; H). Values represent mean±SD (n = 3–4); different letters above the symbols indicate statistically significant differences within each parameter (p < 0.05). Fluorescence traces in panel A are normalized between F_O_ and F_M_ (initial and maximal fluorescence, respectively) and are presented without error bars for clarity. Raw OJIP curves are presented in Supplementary Fig. S12.

The F_O_ and F_M_ values determined during the OJIP transients exhibited similar trends to those derived from the slow fluorescence kinetics (Supplementary Fig. S10). The parameter φP_0_, describing the maximum quantum yield of PSII (Eq. (17)), reached its highest values under 435 nm, 465 nm, and 687 nm cultivation light (Fig. 6B). Rather than indicating intrinsically more efficient PSII photochemistry, this most likely reflects the high PSII/PSI ratio under these wavelengths (Fig. 4F): since φP_0_ was derived from whole-cell F_O_ and F_M_ recorded at 625 nm, the comparatively constant PSI fluorescence contributes a larger relative offset to F_M_ in the low-PSII cultures (495–663 nm), depressing their apparent φP_0_, whereas in the high-PSII cultures the PSII signal dominates (Remelli and Santabarbara, 2018). This same PSII/PSI dependence propagates to the parameters that require φP_0_ for their determination, such as the apparent antenna size of PSII (ABS/RC; Eq. (21)) and the flux of energy dissipated in processes other than trapping per active PSII (DI_0_/RC; Eq. (23)). Both parameters exhibited their lowest values under 435 nm, 465 nm, and 687 nm cultivation light. ABS/RC was the highest in cultures cultivated under red light (Fig. 6C), likely due to the high PC content (Fig. 4E), which secured efficient capturing of the 625 nm ML used for the OJIP transient recording. The combination of efficient ML capturing and low PSII content provides a basis for the high DI_0_/RC under red cultivation light (Fig. 6H), as both parameters are expressed ‘per active PSII’.

Other parameters do not require φP_0_ for their determination and are therefore insensitive to the PSII/PSI-ratio bias described above; these include the efficiency with which a PSII-trapped electron is transferred from Q_A_^-^ to PQ (ψE_0_; Eq. (18)) or from PQH_2_ to the final PSI acceptors (δR_0_; Eq. (19)), as well as the relative variable fluorescence yield at the J-step (V_J_; Eq. (15)), which is related to the redox state of the PQ pool (Tóth et al., 2007; Tsimilli-Michael et al., 2009). The V_J_ parameter reached its highest values in the cultures cultivated under red light (Fig. 6F). This elevated V_J_, together with the most pronounced dip in the fluorescence signal between F_J_ (2 ms) and F_I_ (30 ms; Fig. 6A), suggests a highly reduced PQ pool in these cultures (Zavřel et al., 2026). When the PQ pool gets reduced, the electron transfer from Q_A_ ^-^ to PQ (ψE_0_ ) slows down (Zavřel et al., 2024), as also confirmed here (Fig. 6E). On the other hand, the parameter δR_0_ remained high under red cultivation light, indicating that transfer from PQH_2_ to the PSI acceptors was not itself limiting; the constraint on electron transport under red light therefore lied at Q ^⁻^→PQ step, as reflected by the low ψE_0_. We note again, however, that the OJIP transients were recorded under 625 nm ML, which could lead to an underestimation of the above-mentioned parameters in cultures with high PE content (such as those grown under 520 nm or 555 nm cultivation light). This was further confirmed by the parameter TR_0_/RC (Eq. (22)), representing the maximum trapped exciton flux per active PSII, which was the highest under 663–687 nm cultivation lights, again in line with high PC continent (Fig. 4E).

## 4. Discussion

In this work, light quality acclimation in the CA3-type cyanobacterium *Nostoc* sp. CCAP 1453/38 was characterized across the full PAR spectrum. To our knowledge, this represents the first comprehensive PAR characterization of a CA3 strain. Previous studies on CA3-type cyanobacteria, predominantly focused on *Fremyella diplosiphon*, paid special attention to the molecular regulation of pigment switching under green versus red light and the associated RcaEFC signaling cascade. By integrating our data with parallel datasets on CA1 and non-CA strains obtained under identical conditions, we identify both the canonical CA3 advantage in the green spectral range and novel acclimation features that operate outside the spectral region traditionally associated with CA3.

The CA3-specific PE upregulation provided a clear fitness advantage of *Nostoc* under green light (495–555 nm LEDs) compared with PC-rich strains such as *Cyanobium gracile* or *Synechocystis* sp. PCC 6803, while showing similar trends to the PE-rich strain *Cyanobium* sp. NIVA-CYA 375 (Bernát et al., 2021; Zavřel et al., 2024; Kis et al., 2026). Because all four strains were grown under identical narrow-band LEDs, the relative growth rate enhancement observed during the spectral shift from the 465 nm to 495 nm cultivation light can be compared directly. This increase was 9% in *Synechocystis* (CA1, PC-rich), 33% in *Cyanobium gracile* (non-CA, PC-rich), 74% in *Cyanobium* sp. NIVA-CYA 375 (non-CA, PE-rich), and 168% in *Nostoc* (CA3) (Fig. 2D). An analogous trend was observed under 520 nm and 555 nm light: in comparison with the 465 nm light, the growth rate was enhanced the most also in *Nostoc* relative to the other strains. Importantly, the substantially smaller response in the PE-rich but non-CA *Cyanobium* sp. NIVA-CYA 375 (74% *vs*. 168%) demonstrates that constitutive PE possession alone does not yield a comparable growth advantage, and the dynamic, regulated upregulation enabled by CA3 is required. These results confirm the importance of chromatic acclimation in broadening the spectrum of captured light wavelengths, thereby providing a clear advantage in otherwise growth-limiting spectral niches (Holtrop et al., 2021; Mattei et al., 2025). However, as recognized recently and further confirmed here, additional factors also play a significant role in optimizing light harvesting, such as the PSII:PSI ratio (Kis et al., 2026).

Blue light (465 nm) was generally the least favorable for the growth of *Nostoc*, as well as of *Synechocystis* and both *Cyanobium* strains. This stems from a common cause: less efficient light harvesting by Chl *a* than by PBS. Even though Chl *a* absorbs blue light efficiently, the absorption cross-section of PSII and PSI *via* their internal core antennas (CP43/CP47 and PsaA/PsaB, respectively) is much smaller compared with PBS (You et al., 2023; Zhang et al., 2024). Phycobilisomes thus improve light harvesting in cyanobacteria, but only under favorable wavelengths. Since none of the aforementioned strains contained PUB chromophore that would absorb blue light efficiently, the composition of their PBS, containing PC, APC, and PE, predetermined blue light as the least favorable wavelength for growth. Crucially, blue photons are captured predominantly by PSI, which was typically several-fold more abundant than PSII in *Nostoc* (Fig. 4F). The under-excitation of PSII relative to PSI limits oxygen evolution and the linear electron transport rate (Luimstra et al., 2018), which subsequently constrains NADPH production, carbon fixation, and ultimately cell division (Rodrigues et al., 2023; Höper et al., 2024). This excitonic imbalance was reflected by the elevated Y(ND) in *Nostoc* (Fig. 2G) and the highest PSII content under blue light (Supplementary Fig. S5), consistent with the transcriptional response observed in *Synechocystis* (Luimstra et al., 2020). Consistent with a limitation in excitation delivery rather than in PSII photochemistry as such, the maximum PSII quantum yield remained high under blue light (Fig. 6B) while the functional PSII cross-section at 625 nm was low (Fig. 6C). To overcome this imbalance, cyanobacteria can activate several molecular mechanisms: increasing the total amount of PBS or PSII, or functionally attaching PBS to PSII. Since protein synthesis requires substantial amounts of both energy and reducing equivalents (Burnap, 2015; Faizi et al., 2018), the upregulation of PBS and PSII synthesis further decreases the amount of energy available for cell division. While the resulting ATP/NADPH demand is generally maintained within a stable range across light intensities (Höper et al. 2024), the combined cost of inefficient blue light harvesting and the metabolic burden of PBS and PSII upregulation cannot be compensated by attaching more PBS to PSII.

The slightly increased growth rate observed under violet light (435 nm) compared with blue light (465 nm) was likely a result of increased PUR, which was related mainly to Chl *a* absorption. This PUR increase was identical in *Nostoc* and all aforementioned strains. However, even though Chl *a* was more abundant in PSI than in PSII, the higher PUR stimulated CEF-PSI much less in *Nostoc* than in *Synechocystis* (Zavřel et al., 2024). In *Nostoc*, the higher specific growth rate under 435 nm cultivation light, compared to the 465 nm one was thus likely related to the combination of only a slightly increased CEF-PSI and a shifted macromolecular composition of *Nostoc* biomass (Fig. 3E–H), which showed different trends from those previously reported in *Synechocystis* (Zavřel et al., 2024). Indeed, in strains with low PSI content, such as *Cyanobium* sp. NIVA-CYA 375, a higher PUR under 435 nm LED can lead to ETR(II) stimulation when compared with 465 nm light, as more light is absorbed by PSII, relative to PSI (Kis et al., 2026). Taken together, these results demonstrate a variety of distinct mechanisms and physiological interplays that cyanobacteria utilize to fine-tune light harvesting in diverse spectral niches.

*Nostoc* grew optimally under 555–687 nm light. This wavelength range was also optimal for all aforementioned strains. For the PC-rich strains, such as *Synechocystis* and *Cyanobium gracile*, the optimal growth wavelength was shifted toward red light (596–633 nm), which reflected efficient light absorption by PC. For the PE-rich strain *Cyanobium* sp. NIVA-CYA 375, interestingly, the optimal growth light was not green but rather near far-red (687 nm), which was, however, again related to the low PSI content. As discussed above, broadening the range of optimal growth wavelengths toward green light in *Nostoc* is a direct consequence of the elevated PE content.

Interestingly, the specific growth rate of *Nostoc* did not decrease under near far-red light (687 nm) compared with red light. This result differed from that observed in both *Synechocystis* and *Cyanobium gracile* (Bernát et al., 2021; Zavřel et al., 2024). Under 687 nm light, PUR was increased compared with the 663 nm light (Fig. 2C); however, this was again related to the efficient light absorption by Chl *a* rather than by PBS, as reflected by a sharp reduction in ETR(II) (Fig. 2E). We note, however, that ETR(II) is calculated from σ_II_ and the effective and maximum PSII quantum yields, Y(II) and φP_0_ (Eqs. (10-12)). Of these, σ_II_ was the lowest under violet, blue, and near far-red light (Supplementary Fig. S13), mirroring the low ABS/RC under the same wavelengths (Fig. 6C) — consistent with a small functional PSII cross-section arising from the shifted PSII/PSI ratio and, particularly under blue light, from inefficient capture of the 625 nm ML by PE-rich phycobilisomes. The φP_0_ was, by contrast, elevated under these wavelengths (Fig. 6B). The reduced ETR(II) under near far-red light thus originated primarily from the low σ_II_ probed at 625 nm, rather than from reduced maximum PSII quantum yield. Nevertheless, as shown previously, some strains, such as *Synechocystis*, upregulate their stress response under near far-red light, including transcription of high-light-inducible proteins (Hübschmann et al., 2005). The similar growth rates observed under red and near far-red light in *Nostoc* suggest that such a stress response was not activated in this strain. On the other hand, under the 687 nm light, *Nostoc* exhibited its lowest cellular content of Chl *a* and carotenoids (Fig. 3B, C), together with low PBS abundance and generally the smallest cell size. These results suggest that the downregulation of PBS synthesis compensated for the reduced efficiency of near far-red light capture, as well as for simultaneous PSII upregulation.

The comparable growth rates under green and red light, despite lower PUR under red wavelengths, suggest differential energy transfer efficiencies between PE-rich and PC-rich PBS configurations. While excitation energy transfer through phycobilisomes is known to approach near-100% efficiency from PE, through PC, to APC under optimal conditions (Zilinskas and Greenwald, 1986), recent ultrafast spectroscopy in *Fremyella diplosiphon* PBS revealed that energy transfer from PE disks to PC along the rods occurs in <600 fs, with the charge-transfer character of bilins in the APC core creating a kinetic bottleneck on >10 ps timescales (Sil et al., 2022). The longer transfer pathway through PE-containing rods thus introduces additional opportunities for energy dissipation through PE fluorescence, consistent with the ∼580 nm emission peak in our 77K spectra (Fig. 4O). In PC-dominated rods assembled under red light, this PE-mediated loss is bypassed; furthermore, red light is efficiently captured by APC in the core and by Chl *a* in both photosystems, providing parallel light-harvesting pathways that compensate for the lower PUR. In addition, aromatic residues of linker proteins fine-tune bilin energy states to establish unidirectional energy transfer (Ma et al., 2020). However, the PE→PC transfer in PE-containing rods adds an interface where excitation can be dissipated as fluorescence or heat. Under red light, PC-dominated rods bypass this step, which may further contribute to comparable growth rates under green and red light despite different PUR. We note, however, that pathway-specific transfer efficiencies in *Nostoc* were not assessed directly in this work, and this interpretation requires further measurements.

Interestingly, PE remained the dominant phycobiliprotein in *Nostoc* not only under green light but across the entire 435–555 nm range (Fig. 4E). The CA3 photosensor RcaE has been biochemically characterized as a green/red photoswitch, with its Pg and Pr states absorbing maximally near 535 nm and 672 nm, respectively (Hirose et al., 2013; Nagae et al., 2021). The CA3 response *in vivo*, however, has so far only been tested under green versus red light (Singh and Montgomery, 2011; Pattanaik et al., 2014), and the behavior of CA3 strains under violet, blue, yellow, or near far-red light has not been previously reported to our knowledge. Our data reveal that *Nostoc* maintains a high PE content not only at 555 nm but down to 435 nm, well outside the conventional “green light” range. The simplest interpretation is that wavelengths shorter than ∼500 nm are absorbed too weakly by either Pg or Pr to drive net photoconversion, leaving the photosensor in or near its default Pg state and PE genes activated. The CA3 fitness advantage in spectral niches dominated by PE absorption may therefore extend considerably beyond the range implied by RcaE’s nominal “green/red” classification—a point with direct ecological relevance for shallow turbid waters, where both blue and green photons may dominate.

Our data address key questions highlighted in recent reviews: the need to integrate the molecular mechanisms of CA3 with their physiological consequences across diverse cyanobacterial taxa, and to incorporate physiological context for a better understanding of phenotypic plasticity in chromatic acclimators (Mondal et al., 2024; Kehoe et al., 2025). By combining genome analysis, which confirms a complete CA3 gene set in *Nostoc* sp. CCAP 1453/38, with full-PAR physiology, the present work establishes a trajectory toward closing both of these gaps.

## 5. Conclusions

This study provides the first comprehensive full-PAR characterization of a CA3-type cyanobacterium, *Nostoc* sp. CCAP 1453/38, and reveals the energetic trade-offs underlying its spectral acclimation. While CA3-driven adjustment of PC and PE provides a clear fitness advantage under green and red light by matching antenna composition to spectral availability, the response of *Nostoc* to wavelengths poorly absorbed by PBS reveals an equally important dimension of spectral acclimation. Under blue light that constrains growth, *Nostoc* attempts to compensate for low PSII excitation by upregulating total PBS and PSII levels and biasing phycobilisome coupling toward PSII (an increased PBS–PSII/PBS–PSI ratio); yet such a metabolically expensive response cannot resolve the underlying excitonic imbalance between PSII and PSI. Under near far-red light, in contrast, *Nostoc* enhances PBS–PSII coupling itself without the metabolic burden of PBS upregulation, which allows it to maintain a specific growth rate as high as that observed under green and red light. These qualitatively different physiological responses to the same challenge of poor PBS absorption and PSII under-excitation expand our understanding of spectral acclimation strategies beyond the CA3 framework and outside the CA-favored spectral range. Together with parallel datasets on CA1 and non-CA strains, our findings clarify how excitation balance and energy allocation limit performance under specific wavelengths in a CA3 strain, extend CA3 characterization beyond *F. diplosiphon*, and provide a framework for optimizing light spectra in controlled cyanobacterial cultivations (Rodrigues et al., 2023), where avoiding metabolically costly acclimation states is as important as choosing wavelengths well-absorbed by PBS.

## Supporting information

Supplementary information with Supplementary Figures S1-S13

## 6. Supplementary Data

## 7. Data Availability

All data and materials are deposited in the online repository available at https://doi.org/10.6084/m9.figshare.33071699. Additionally, a set of freely accessible tools for the streamlined processing and analysis of Chl *a* fluorescence and spectrofluorometry data, alongside a set of image processing tools used for the quantification of cell diameter and pigment-specific fluorescence signal intensities, is provided at https://www.cyano.tools/.

## 8. Funding

This work was funded by the Ministry of Education, Youth and Sports of the Czech Republic (LUAUS24149) and by the National Multidisciplinary Laboratory for Climate Change (RRF-2.3.1-21-2022-00014) project within the framework of Hungary’s National Recovery and Resilience Plan supported by the Recovery and Resilience Facility of the European Union.

## 9. Acknowledgments

## 10. Author Contributions

TZ and GB designed the experiments. TZ and AP performed the experiments and analysed the results. ML performed FTIR measurements and analysed the data. LK performed the HPLC measurements and analysed the data. ZN performed the confocal microscopy evaluations. MK recorded the 77K fluorescence spectra and analyzed the data. JM provided the bioinformatics analysis. AS performed the statistical analysis. TZ wrote the manuscript. All authors reviewed, revised, and approved the final manuscript for publication.

## Disclosures

The authors have no conflicts of interest to declare.

## Notes

https://doi.org/10.6084/m9.figshare.33071699

### Competing Interest Statement

The authors have declared no competing interest.

## References

Bernát, G., Steinbach, G., Kaňa, R., Govindjee, Misra, A. N., and Prašil, O. (2018). On the origin of the slow M-T chlorophyll a fluorescence decline in cyanobacteria: interplay of short-term light-responses. Photosynth Res 136, 183–198.

Bernát, G., Zavřel, T., Kotabová, E., Kovács, L., Steinbach, G., Vörös, L., et al. (2021). Photomorphogenesis in the Picocyanobacterium Includes Increased Phycobilisome Abundance Under Blue Light, Phycobilisome Decoupling Under Near Far-Red Light, and Wavelength-Specific Photoprotective Strategies. Front Plant Sci 12, 612302.

Burnap, R. L. (2015). Systems and Photosystems: Cellular Limits of Autotrophic Productivity in Cyanobacteria. Front. Bioeng. Biotechnol. 3, 117998.

Calzadilla, P. I., and Kirilovsky, D. (2020). Revisiting cyanobacterial state transitions. Photochem Photobiol Sci 19, 585–603.

Carvalho, M. C., Schulz, K. G., and Eyre, B. D. (2017). Respiration of new and old carbon in the surface ocean: Implications for estimates of global oceanic gross primary productivity. Global Biogeochem. Cycles 31, 975–984.

Chang, L., Liu, X., Li, Y., Liu, C.-C., Yang, F., Zhao, J., et al. (2015). Structural organization of an intact phycobilisome and its association with photosystem II. Cell Res 25, 726–737.

Chmelík, D., Hrouzek, P., Fedorko, J., Vu, D. L., Urajová, P., Mareš, J., et al. (2019). Accumulation of cyanobacterial oxadiazine nocuolin A is enhanced by temperature shift during cultivation and is promoted by bacterial co-habitants in the culture. Algal Res. 44, 101673.

Cloern, J. E. (1987). Turbidity as a control on phytoplankton biomass and productivity in estuaries. Cont. Shelf Res. 7, 1367–1381.

Delcher, A. L., Bratke, K. A., Powers, E. C., and Salzberg, S. L. (2007). Identifying bacterial genes and endosymbiont DNA with Glimmer. Bioinformatics 23, 673–679.

Dufour, L., Garczarek, L., Mattei, F., Gouriou, B., Clairet, J., Ratin, M., et al. (2025). Competition for light color between marine strains with fixed and variable pigmentation. *Appl Environ Microbiol*, e0008725.

Faizi, M., Zavřel, T., Loureiro, C., Červený, J., and Steuer, R. (2018). A model of optimal protein allocation during phototrophic growth. Biosystems 166, 26–36.

Felcmanová, K., Lukeš, M., Kotabová, E., Lawrenz, E., Halsey, K. H., and Prášil, O. (2017). Carbon use efficiencies and allocation strategies in Prochlorococcus marinus strain PCC 9511 during nitrogen-limited growth. Photosynth Res 134, 71–82.

Flombaum, P., Gallegos, J. L., Gordillo, R. A., Rincón, J., Zabala, L. L., Jiao, N., et al. (2013). Present and future global distributions of the marine Cyanobacteria Prochlorococcus and Synechococcus. Proc Natl Acad Sci U S A 110, 9824–9829.

Fox, J., and Weisberg, S. (2011). An R Companion to Applied Regression. SAGE Publications.

Giraudoux, P., Antonietti, J.-P., Beale, C., Groemping, U., Lancelot, R., Pleydell, D., et al. (2023). pgirmess: spatial analysis and data mining for field ecologists. Available at: https://cran.r-project.org/web/packages/pgirmess/pgirmess.pdf

Hirose, Y., Chihong, S., Watanabe, M., Yonekawa, C., Murata, K., Ikeuchi, M., et al. (2019). Diverse Chromatic Acclimation Processes Regulating Phycoerythrocyanin and Rod-Shaped Phycobilisome in Cyanobacteria. Mol Plant 12, 715–725.

Hirose, Y., Rockwell, N. C., Nishiyama, K., Narikawa, R., Ukaji, Y., Inomata, K., et al. (2013). Green/red cyanobacteriochromes regulate complementary chromatic acclimation via a protochromic photocycle. Proc Natl Acad Sci U S A 110, 4974–4979.

Holtrop, T., Huisman, J., Stomp, M., Biersteker, L., Aerts, J., Grébert, T., et al. (2021). Vibrational modes of water predict spectral niches for photosynthesis in lakes and oceans. Nat Ecol Evol 5, 55–66.

Höper, R., Komkova, D., Zavřel, T., and Steuer, R. (2024). A quantitative description of light-limited cyanobacterial growth using flux balance analysis. PLoS Comput Biol 20, e1012280.

Hrouzek, P., Kapuścik, A., Vacek, J., Voráčová, K., Paichlová, J., Kosina, P., et al. (2016). Cytotoxicity evaluation of large cyanobacterial strain set using selected human and murine in vitro cell models. Ecotoxicol Environ Saf 124, 177–185.

Hübschmann, T., Yamamoto, H., Gieler, T., Murata, N., and Börner, T. (2005). Red and far-red light alter the transcript profile in the cyanobacterium Synechocystis sp. PCC 6803: impact of cyanobacterial phytochromes. FEBS Lett 579, 1613–1618.

Hyun, B., Kim, J.-M., Jang, P.-G., Jang, M.-C., Choi, K.-H., Lee, K., et al. (2020). The effects of ocean acidification and warming on growth of a natural community of coastal phytoplankton. J. Mar. Sci. Eng. 8, 821.

Kehoe, D. M., Biswas, A., Chen, B., Dufour, L., Grébert, T., Haney, A. M., et al. (2025). Light Color Regulation of Photosynthetic Antennae Biogenesis in Marine Phytoplankton. Plant Cell Physiol 66, 168–180.

Kerfeld, C. A., Melnicki, M. R., Sutter, M., and Dominguez-Martin, M. A. (2017). Structure, function and evolution of the cyanobacterial orange carotenoid protein and its homologs. New Phytol 215, 937–951.

Kirilovsky, D., and Kerfeld, C. A. (2016). Cyanobacterial photoprotection by the orange carotenoid protein. Nat Plants 2, 16180.

Kis, M., Zavřel, T., Fodor, I., Segečová, A., Urbán, P., Gálik, B., et al. (2026). Low PSI content broadens the optimal light spectrum for a phycoerythrin-dominated cyanobacterium towards near far-red. Sci Rep 16. doi: 10.1038/s41598-026-50772-z

Liu, H., Lu, Y., Wolf, B., Saer, R., King, J. D., and Blankenship, R. E. (2018). Photoactivation and relaxation studies on the cyanobacterial orange carotenoid protein in the presence of copper ion. Photosynth Res 135, 143–147.

Luimstra, V. M., Schuurmans, J. M., Hellingwerf, K. J., Matthijs, H. C. P., and Huisman, J. (2020). Blue light induces major changes in the gene expression profile of the cyanobacterium Synechocystis sp. PCC 6803. Physiol Plant 170, 10–26.

Luimstra, V. M., Schuurmans, J. M., Verschoor, A. M., Hellingwerf, K. J., Huisman, J., and Matthijs, H. C. P. (2018). Blue light reduces photosynthetic efficiency of cyanobacteria through an imbalance between photosystems I and II. Photosynth Res 138, 177–189.

Ma, J., You, X., Sun, S., Wang, X., Qin, S., and Sui, S.-F. (2020). Structural basis of energy transfer in Porphyridium purpureum phycobilisome. Nature 579, 146–151.

Mareš, J., Strunecký, O., Bučinská, L., and Wiedermannová, J. (2019). Evolutionary Patterns of Thylakoid Architecture in Cyanobacteria. Front Microbiol 10, 277.

Martiny, A. C., Pham, C. T. A., Primeau, F. W., Vrugt, J. A., Moore, J. K., Levin, S. A., et al. (2013). Strong latitudinal patterns in the elemental ratios of marine plankton and organic matter. Nat. Geosci. 6, 279–283.

Matsumoto, K., Guo, Z., and Maas, A. E. (2026). Stoichiometric modulation of zooplankton grazing on ocean organic matter biogeochemistry: Results from idealized food web modeling. Ecol. Modell. 513, 111423.

Mattei, F., Hickman, A. E., Uitz, J., Dufour, L., Vellucci, V., Garczarek, L., et al. (2025). Chromatic acclimation shapes phytoplankton biogeography. Sci Adv 11, eadr9609.

Mattei, F., and Scardi, M. (2021). Collection and analysis of a global marine phytoplankton primary-production dataset. Earth Syst. Sci. Data 13, 4967–4985.

Mondal, S., Pandey, D., and Singh, S. P. (2024). Chromatic acclimation in cyanobacteria renders robust photosynthesis and fitness in dynamic light environment: Recent advances and future perspectives. Physiol Plant 176, e14536.

Müller, S., Zavřel, T., and Červený, J. (2019). Towards a quantitative assessment of inorganic carbon cycling in photosynthetic microorganisms. Eng Life Sci 19, 955–967.

Nagae, T., Unno, M., Koizumi, T., Miyanoiri, Y., Fujisawa, T., Masui, K., et al. (2021). Structural basis of the protochromic green/red photocycle of the chromatic acclimation sensor RcaE. Proc Natl Acad Sci U S A 118. doi: 10.1073/pnas.2024583118

Palenik, B. (2001). Chromatic adaptation in marine Synechococcus strains. Appl Environ Microbiol 67, 991–994.

Pattanaik, B., Busch, A. W. U., Hu, P., Chen, J., and Montgomery, B. L. (2014). Responses to iron limitation are impacted by light quality and regulated by RcaE in the chromatically acclimating cyanobacterium Fremyella diplosiphon. Microbiology (Reading) 160, 992–1005.

Pfennig, T., Kullmann, E., Zavřel, T., Nakielski, A., Ebenhöh, O., Červený, J., et al. (2024). Shedding light on blue-green photosynthesis: A wavelength-dependent mathematical model of photosynthesis in Synechocystis sp. PCC 6803. PLoS Comput Biol 20, e1012445.

Rabouille, S., Campbell, D. A., Masuda, T., Zavřel, T., Bernát, G., Polerecky, L., et al. (2021). Electron & Biomass Dynamics of Under Interacting Nitrogen & Carbon Limitations. Front Microbiol 12, 617802.

R Core Team (2022). R: A language and environment for statistical computing. Available at: https://www.r-project.org/

Remelli, W., and Santabarbara, S. (2018). Excitation and emission wavelength dependence of fluorescence spectra in whole cells of the cyanobacterium Synechocystis sp. PPC6803: Influence on the estimation of Photosystem II maximal quantum efficiency. Biochim Biophys Acta Bioenerg 1859, 1207–1222.

Righetti, D., Vogt, M., Gruber, N., Psomas, A., and Zimmermann, N. E. (2019). Global pattern of phytoplankton diversity driven by temperature and environmental variability. Sci Adv 5, eaau6253.

Rippka, R., Stanier, R. Y., Deruelles, J., Herdman, M., and Waterbury, J. B. (1979). Generic assignments, strain histories and properties of pure cultures of Cyanobacteria. Microbiology 111, 1–61.

Rodrigues, J. S., Kovács, L., Lukeš, M., Höper, R., Steuer, R., Červený, J., et al. (2023). Characterizing isoprene production in cyanobacteria - Insights into the effects of light, temperature, and isoprene on Synechocystis sp. PCC 6803. Bioresour Technol 380, 129068.

Sanfilippo, J. E., Garczarek, L., Partensky, F., and Kehoe, D. M. (2019). Chromatic Acclimation in Cyanobacteria: A Diverse and Widespread Process for Optimizing Photosynthesis. Annu Rev Microbiol 73, 407–433.

Scheer, H., and Zhao, K.-H. (2008). Biliprotein maturation: the chromophore attachment. Mol Microbiol 68, 263–276.

Schreiber, U., Klughammer, C., and Kolbowski, J. (2012). Assessment of wavelength-dependent parameters of photosynthetic electron transport with a new type of multi-color PAM chlorophyll fluorometer. Photosynth Res 113, 127–144.

Sil, S., Tilluck, R. W., Mohan T M, N., Leslie, C. H., Rose, J. B., Domínguez-Martín, M. A., et al. (2022). Excitation energy transfer and vibronic coherence in intact phycobilisomes. Nat Chem 14, 1286–1294.

Singh, S. P., and Montgomery, B. L. (2011). Determining cell shape: adaptive regulation of cyanobacterial cellular differentiation and morphology. Trends Microbiol 19, 278–285.

Stirbet, A., Lazár, D., Kromdijk, J., and Govindjee, G. (2018). Chlorophyll a fluorescence induction: Can just a one-second measurement be used to quantify abiotic stress responses? Photosynthetica 56, 86–104.

Stirbet, A., Lazár, D., Papageorgiou, G. C., and Govindjee (2019). “Chlorophyll a fluorescence in Cyanobacteria: Relation to photosynthesis,” in Cyanobacteria, (Elsevier), 79–130.

Stomp, M., Huisman, J., Stal, L. J., and Matthijs, H. C. P. (2007). Colorful niches of phototrophic microorganisms shaped by vibrations of the water molecule. ISME J 1, 271–282.

Sugie, K., Fujiwara, A., Nishino, S., Kameyama, S., and Harada, N. (2020). Impacts of temperature, CO2, and salinity on phytoplankton community composition in the western arctic ocean. Front. Mar. Sci. 6. doi: 10.3389/fmars.2019.00821

Tamary, E., Kiss, V., Nevo, R., Adam, Z., Bernát, G., Rexroth, S., et al. (2012). Structural and functional alterations of cyanobacterial phycobilisomes induced by high-light stress. Biochim Biophys Acta 1817, 319–327.

Tchernov, D., Silverman, J., Luz, B., Reinhold, L., and Kaplan, A. (2003). Massive light-dependent cycling of inorganic carbon between oxygenic photosynthetic microorganisms and their surroundings. Photosynth Res 77, 95–103.

Tóth, S. Z., Schansker, G., and Strasser, R. J. (2007). A non-invasive assay of the plastoquinone pool redox state based on the OJIP-transient. Photosynth Res 93, 193–203.

Tsimilli-Michael, M., Stamatakis, K., and Papageorgiou, G. C. (2009). Dark-to-light transition in Synechococcus sp. PCC 7942 cells studied by fluorescence kinetics assesses plastoquinone redox poise in the dark and photosystem II fluorescence component and dynamics during state 2 to state 1 transition. Photosynth Res 99, 243–255.

Tsuyama, M., Shibata, M., Kawazu, T., and Kobayashi, Y. (2004). An Analysis of the Mechanism of the Low-wave Phenomenon of Chlorophyll Fluorescence. Photosynth Res 81, 67–76.

van Amerongen, H., and Croce, R. (2025). Nonphotochemical quenching in plants: Mechanisms and mysteries. Plant Cell 37. doi: 10.1093/plcell/koaf240

van Stokkum, I. H. M., Akhtar, P., Biswas, A., and Lambrev, P. H. (2023). Energy transfer from phycobilisomes to photosystem I at 77 K. Front Plant Sci 14, 1293813.

Voráčová, K., Hájek, J., Mareš, J., Urajová, P., Kuzma, M., Cheel, J., et al. (2017). The cyanobacterial metabolite nocuolin a is a natural oxadiazine that triggers apoptosis in human cancer cells. PLoS One 12, e0172850.

Watanabe, M., and Ikeuchi, M. (2013). Phycobilisome: architecture of a light-harvesting supercomplex. Photosynth Res 116, 265–276.

Wirtz, K., and Smith, S. L. (2021). Author Correction: Vertical migration by bulk phytoplankton sustains biodiversity and nutrient input to the surface ocean. Sci Rep 11, 9486.

You, X., Zhang, X., Cheng, J., Xiao, Y., Ma, J., Sun, S., et al. (2023). In situ structure of the red algal phycobilisome-PSII-PSI-LHC megacomplex. Nature 616, 199–206.

Zavřel, T., Chmelík, D., Sinetova, M. A., and Červený, J. (2018a). Spectrophotometric Determination of Phycobiliprotein Content in Cyanobacterium Synechocystis. J Vis Exp. doi: 10.3791/58076

Zavřel, T., Faizi, M., Loureiro, C., Poschmann, G., Stühler, K., Sinetova, M., et al. (2019). Quantitative insights into the cyanobacterial cell economy. Elife 8. doi: 10.7554/eLife.42508

Zavřel, T., Pohland, A.-C., Pfennig, T., Matuszyńska, A. B., Tóth, S. Z., Bernát, G., et al. (2026). Estimating the redox state of the plastoquinone pool in algae and cyanobacteria via OJIP fluorescence: perspectives and limitations. Photosynth Res 164, 6.

Zavřel, T., Segečová, A., Kovács, L., Lukeš, M., Novák, Z., Pohland, A.-C., et al. (2024). A Comprehensive Study of Light Quality Acclimation in Synechocystis Sp. PCC 6803. Plant Cell Physiol 65, 1285–1297.

Zavřel, T., Szabó, M., Tamburic, B., Evenhuis, C., Kuzhiumparambil, U., Literáková, P., et al. (2018b). Effect of carbon limitation on photosynthetic electron transport in Nannochloropsis oculata. J Photochem Photobiol B 181, 31–43.

Zhang, X., Xiao, Y., You, X., Sun, S., and Sui, S.-F. (2024). In situ structural determination of cyanobacterial phycobilisome-PSII supercomplex by STAgSPA strategy. Nat Commun 15, 7201.

Zilinskas, B. A., and Greenwald, L. S. (1986). Phycobilisome structure and function. Photosynth Res 10, 7–35.

Zitoun, R., Marcinek, S., Hatje, V., Sander, S. G., Völker, C., Sarin, M., et al. (2024). Climate change driven effects on transport, fate and biogeochemistry of trace element contaminants in coastal marine ecosystems. Commun. Earth Environ. 5. doi: 10.1038/s43247-024-01679-y

