## Supplementary information with Supplementary Figures S1-S13 for "Metabolic cost as a determinant of light quality acclimation: a full-PAR characterization of the CA3 cyanobacterium *Nostoc* sp. CCAP 1453/38"

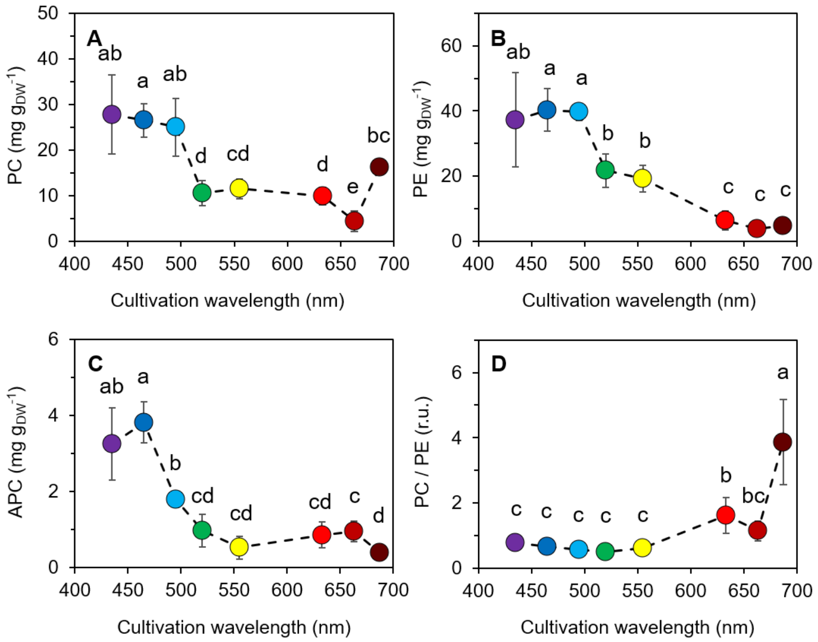


**Figure S1** Content and ratio of phycobiliproteins in *Nostoc* cells. Cellular content of phycocyanin (PC; panel A), phycoerythrin (PE; panel B), and allophycocyanin (APC; panel C) was determined from spectrophotometrically quantified PBS and cellular dry weight (DW) measurements (see Materials and Methods for details). The values represent mean±SD (n = 3–4); the different letters above the symbols indicate statistically significant differences within each parameter ($p < 0.05$). Pigment content is shown relative to the cellular dry weight (DW).

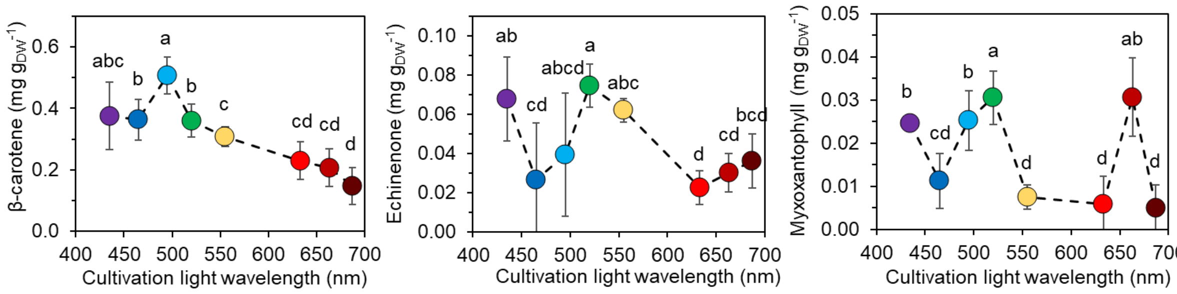


**Figure S2** Content of carotenoids in *Nostoc* cells. Cellular content of β-carotene (A), echinenone (B) and myxoxanthophyll (C) was determined by HPLC analysis and cellular dry weight (DW) measurements (see Material and Methods for details). The values represent mean±SD (n = 3–4); the different letters above the symbols indicate statistically significant differences within each parameter ($p < 0.05$). Carotenoids content is shown relative to the cellular dry weight (DW).

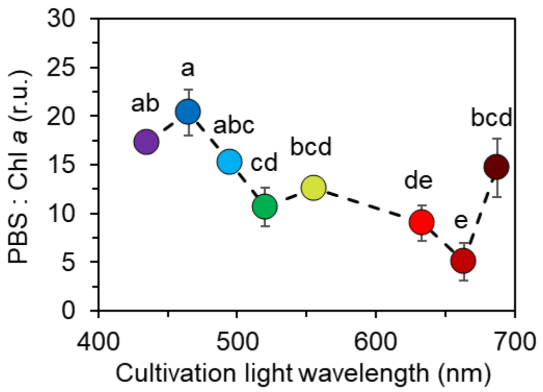


**Figure S3** The ratio of phycobilisomes (PBS) to chlorophyll *a* (Chl *a*) in *Nostoc* cells, as calculated from HPLC analysis and cellular dry weight measurements (see Materials and Methods for details). The values represent mean±SD (n = 3–4); the different letters above the symbols indicate statistically significant differences within each parameter ($p < 0.05$).

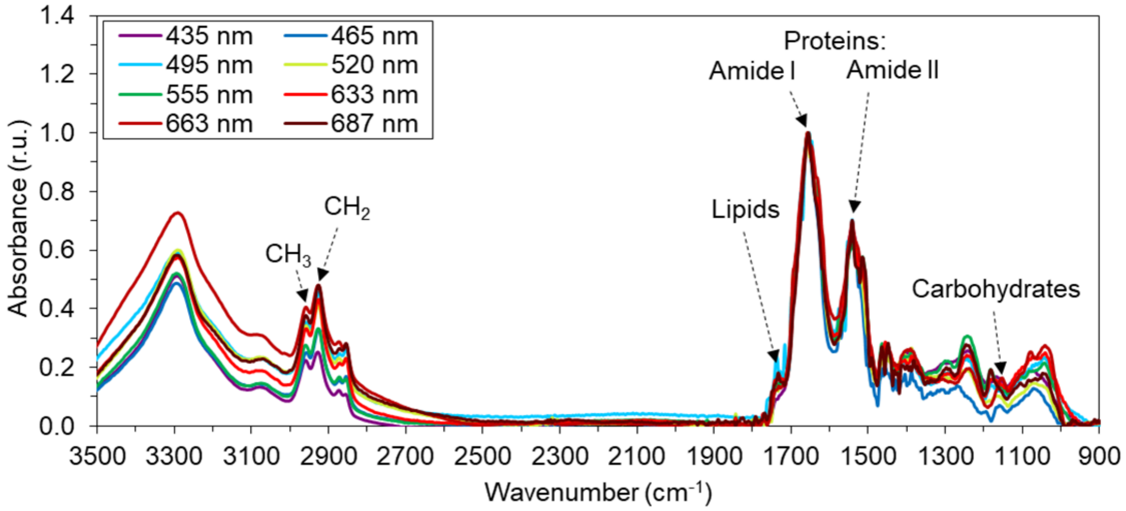


**Figure S4** Representative FTIR spectra of *Nostoc* lyophilized biomass. The spectra were normalized to the Amide I peak (~1652 $cm^{-1}$). Lipids and carbohydrates were evaluated from the amplitude of the peaks at ~1735 $cm^{-1}$ and ~1152 $cm^{-1}$, respectively. The length of fatty acid chain was estimated from a ratio of amplitudes of the asymmetric vibrations of $CH_3$ at ~2960 $cm^{-1}$ and $CH_2$ at ~2925 $cm^{-1}$.

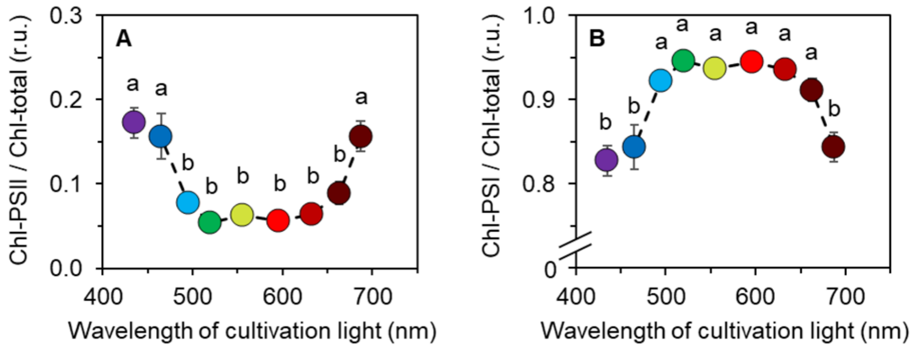


**Figure S5** Estimation of the relative content of PSII (Chl-PSII) and PSI (Chl-PSI) in the total PSII+PSI pool in *Nostoc* cells, based on 77K fluorescence measured upon excitation at 440 nm (exciting primarily Chlorophyll *a*) and emission at 695 nm (PSII) and 726 nm (PSI), (see Eqs. 3 and 4). The values represent mean±SD (n = 3–4); the different letters above the symbols indicate statistically significant differences within each parameter ($p < 0.05$).

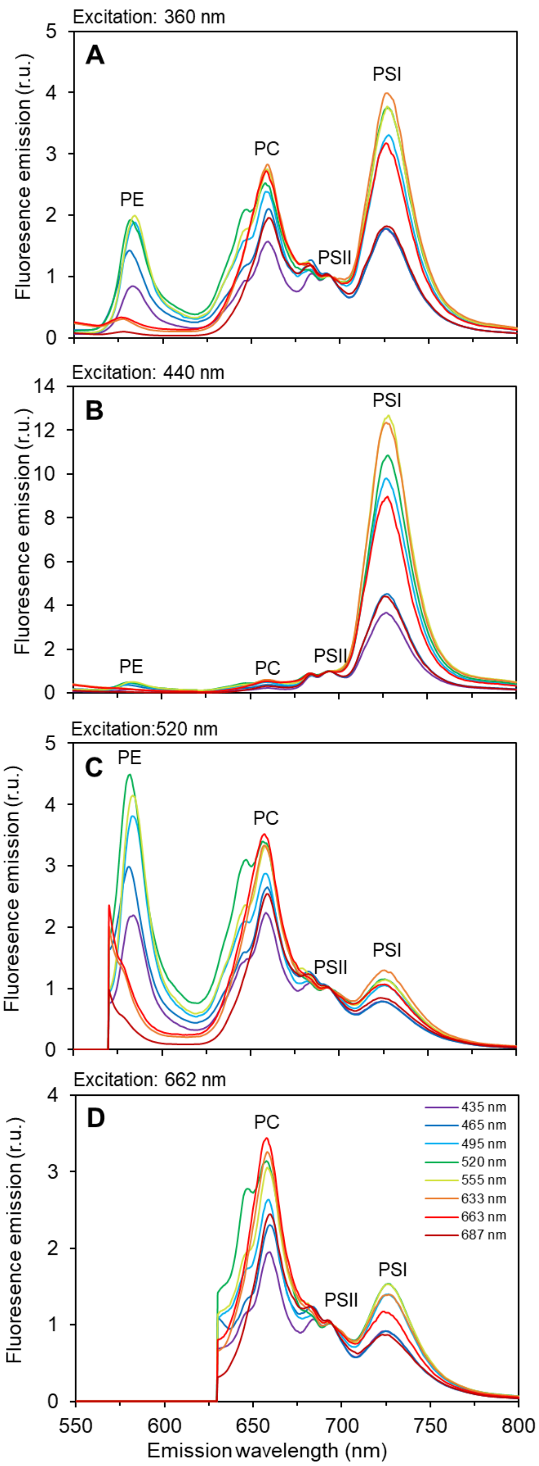


**Figure S6** 77K fluorescence emission spectra of *Nostoc* cultures, recorded upon excitation at 360 nm (A), 440 nm (B), 520 nm (C), and 662 nm (D). The spectra were normalized to the PSII emission peak at 689 nm, and represent averages of 3–4 biological replicates; error intervals are not shown for clarity. The fluorescence emission peaks used in Eqs. (3–9) include phycoerythrin (PE; emission peak ~580 nm), phycocyanin (PC; emission peak ~662 nm), photosystem II (PSII; emission peak ~689 nm), and photosystem I (PSI; emission peak ~724 nm).

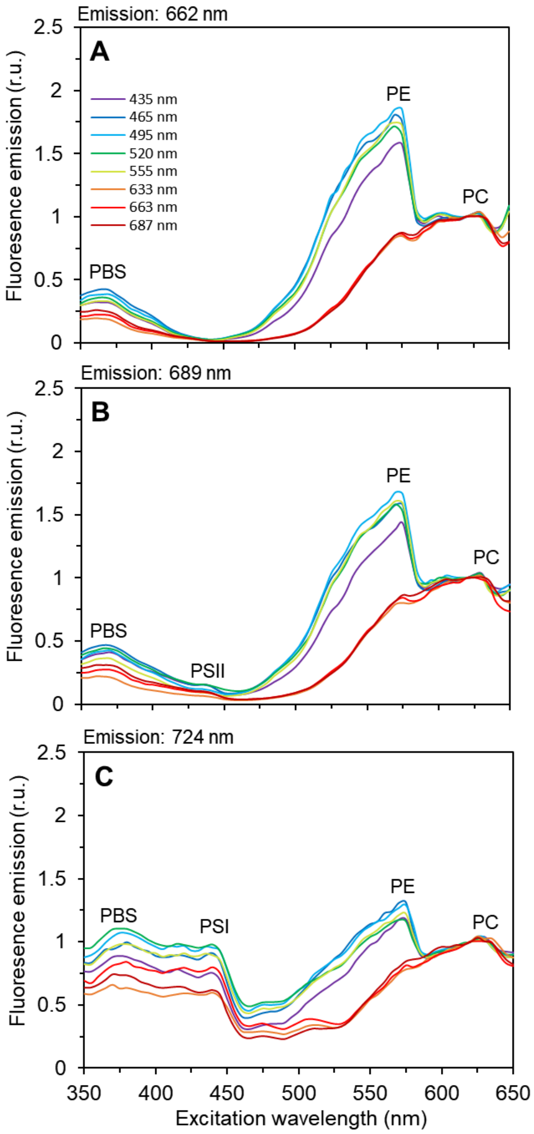


**Figure S7** 77K fluorescence excitation spectra of *Nostoc* cultures, recorded upon emission at 662 nm (phycocyanin, PC; panel A), 689 nm (photosystem II, PSII; panel B) and 724 nm (photosystem I, PSI; panel C). The spectra were normalized to the phycocyanin excitation at 620 nm, and represent averages of 3–4 biological replicates; error intervals are not shown for clarity. The fluorescence excitation peaks used in Eqs. (3–9) include PSI and PSII (excitation at 440 nm), PE (excitation at 560 nm) and PC (excitation at 620 nm).

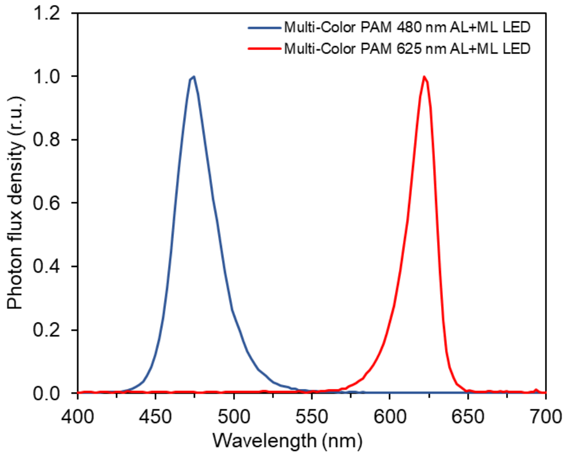


**Figure S8** Emission spectra of blue and red LEDs of the Multi-Color PAM, used as actinic light (AL) and measuring light (ML) source during fast (OJIP curves) and slow fluorescence kinetic measurements (estimating the kinetics of state transition and NPQ). The spectra were recorded using a SpectraPen mini (Photon Systems Instruments, Czechia) and are shown as normalized to their maximal value.

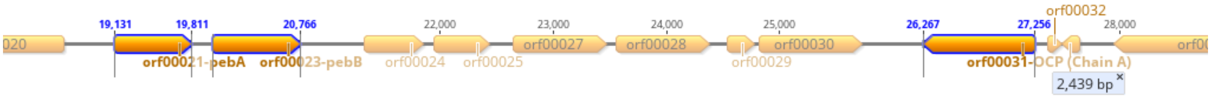


**Figure S9** Identification of orange carotenoid protein (OCP) in the genome of *Nostoc*. The gene encoding OCP was identified by BLASTp and conserved domain analysis in a genomic contig containing also the pebAB operon involved in phycoerythrobilin biosynthesis.

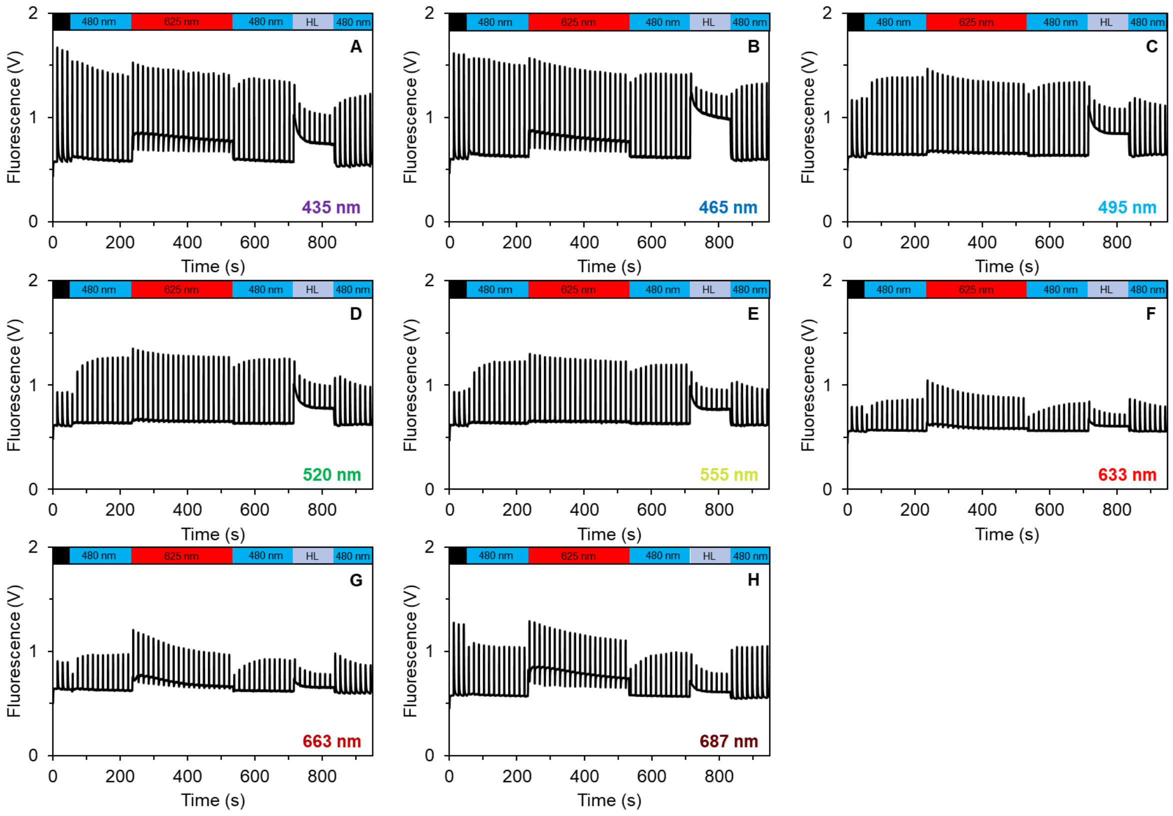


**Figure S10** Chlorophyll *a* fluorescence traces from which the ETR(II), NPQ, and rate of state transition were determined in *Nostoc* cultures cultivated under 435–687 nm lights (panels A–H). The color bars above the charts represent the dark acclimation period (black), 480 nm actinic light (AL) of photon flux density (PFD) 80 μmol photons $m^{-2}$ $s^{-1}$ inducing State I, 625 nm AL of PFD 50 μmol photons $m^{-2}$ $s^{-1}$ inducing State II (Calzadilla and Kirilovsky 2020), or 480 nm AL of PFD 1 800 μmol photons $m^{-2}$ $s^{-1}$ inducing NPQ. The spectra represent averages of 3–4 biological replicates; error intervals are not shown for clarity. We note that the amplitude of $F_M$' under 480 nm AL in MC-PAM was likely underestimated (Zavřel et al. 2024).

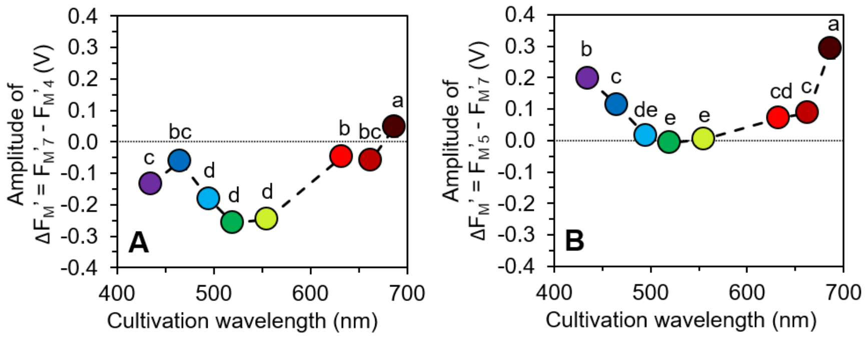


**Figure S11** $F_M$' shift during slow fluorescence induction kinetics. Panel A shows the amplitude of difference between the last $F_M$' point ($F_M$'$_7$ according to Fig. 5) and the $F_M$' measured at the end of the second illumination phase under 480 nm actinic light ($F_M$'$_4$ according to Fig. 5). Panel B shows the amplitude of difference between the $F_M$' measured at the at the end of the high light treatment ($F_M$'$_5$ according to Fig. 5) and the last $F_M$' point ($F_M$'$_7$ according to Fig. 5). The values represent mean±SD (n = 3–4); the different letters above the symbols indicate statistically significant differences within each parameter ($p < 0.05$).

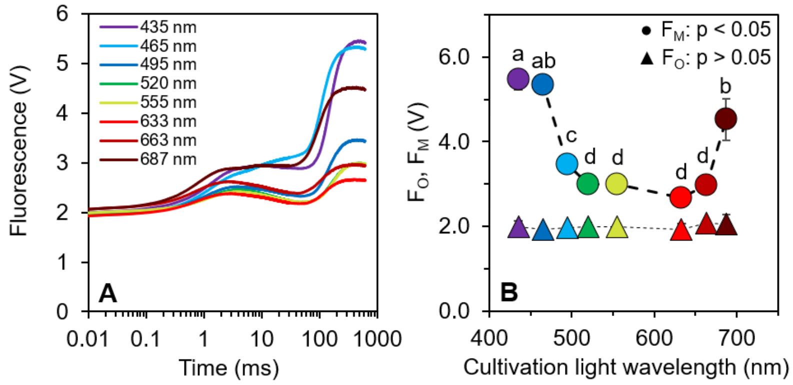


**Figure S12** Raw OJIP curves (A) and derived $F_O$ and $F_M$ parameters (B) of dark-acclimated *Nostoc* cultures. The OJIP curves represent averages of 3–4 biological replicates; error intervals are not shown for clarity. The values in panel B represent mean±SD (n = 3–4); the different letters above the symbols indicate statistically significant differences within each parameter ($p < 0.05$).

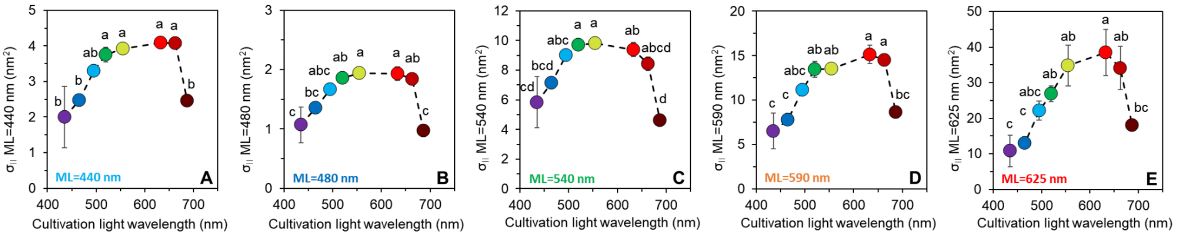

**Figure S13** Functional absorption cross-section of PSII ($\sigma_{II}$) measured under measuring light (ML) at 440 nm (A), 480 nm (B), 540 nm (C), 590 nm (D), and 625 nm (E). The values represent mean±SD (n = 3–4); the different letters above the symbols indicate statistically significant differences within each parameter ($p < 0.05$).

**References:**


Calzadilla, Pablo I., and Diana Kirilovsky. 2020. “Revisiting Cyanobacterial State Transitions.” *Photochemical & Photobiological Sciences : Official Journal of the European Photochemistry Association and the European Society for Photobiology* 19 (5): 585–603.

Zavřel, Tomáš, Anna Segečová, László Kovács, et al. 2024. “A Comprehensive Study of Light Quality Acclimation in Synechocystis Sp. PCC 6803.” *Plant & Cell Physiology* 65 (8): 1285–1297.
